# Comprehensive evaluation of LLM capabilities for interpretation and analysis of genome-scale metabolic models in metabolic engineering

**DOI:** 10.64898/2026.06.03.730004

**Authors:** Jing Wui Yeoh, C. Pawan K. Patro, Limsoon Wong, Chueh Loo Poh

**Affiliations:** National Centre for Engineering Biology (NCEB), 15 Kent Ridge Crescent, Singapore 119276, Singapore; NUS Synthetic Biology for Clinical and Technological Innovation (SynCTI), Singapore 117456, Singapore; School of Computing, National University of Singapore, 13 Computing Drive, Singapore 117417, Singapore; Department of Biomedical Engineering, College of Design and Engineering, National University of Singapore, Singapore 119077, Singapore

**Keywords:** LLM benchmarking, Genome-scale metabolic model, multi-LLM auto-evaluation, pathway/strain engineering, metabolic flux prediction

## Abstract

Genome-scale metabolic models (GSMs) underpin pathway and strain engineering by linking genes to metabolic reactions and enabling system-level simulation of cellular fluxes and intervention effects, yet end-to-end analysis workflows remain fragmented, expert-demanding, and slow to adapt. Large language models (LLMs) could transform this landscape, lowering the barrier by explaining concepts, interpreting GSM files, and turning natural-language instructions into valid analysis code, thereby substantially mitigating the time, effort, and expertise required. However, their reliability for domain-specific tasks remains unexplored. Here, we delivered a systematic benchmark of four leading LLMs (GPT-4, Gemini, Claude, DeepSeek-R1) across four task areas central to metabolic engineering: domain knowledge, metabolic flux prediction, pathway construction, and flux optimization. For benchmarking, we introduced a standardized, rubric-based evaluation framework that uses multi-LLM automated scoring (an ensemble of LLM-as-a-judge assessments) and two distinct sets of nine task-tailored metrics (domain vs coding-focused tasks), rated on a 1-5 scale (up to 45 per task), covering scientific validity and code executability where applicable. Across tasks, we reveal consistent strengths (conceptual explanation, code synthesis) and critical failure modes (e.g., context window limitations, incorrect identifier assumptions, strain-dependent reasoning errors, and errors in domain-specific algorithms). In aggregate, DeepSeek-R1 led in domain tasks, narrowly edging GPT-4, Claude, and Gemini, demonstrating that conceptual biological logic remains highly invariant across architectures. In contrast, Gemini achieved the highest score for coding tasks, distinguished by functional execution and excelled in error handling, documentation, and readability, followed by GPT-4, Claude, and DeepSeek. We also evaluated LLM self-inspection capability by injecting subtle, consequential faults: a stoichiometric sign error causing mass imbalance and an omitted pathway reaction. We reveal that conversational “blind search” prompting completely fails to localize these network faults. Instead, robust error localization requires prompts reframed with domain-informed constraints that force the LLM to leverage tool-assisted code procedures, such as COBRApy mass-balance functions. Together, this work establishes an evidence-based baseline for LLM-enabled GSM analysis, providing actionable guidance for building reliable, automation-ready workflows for pathway and strain design.

**Graphical Abstract:** 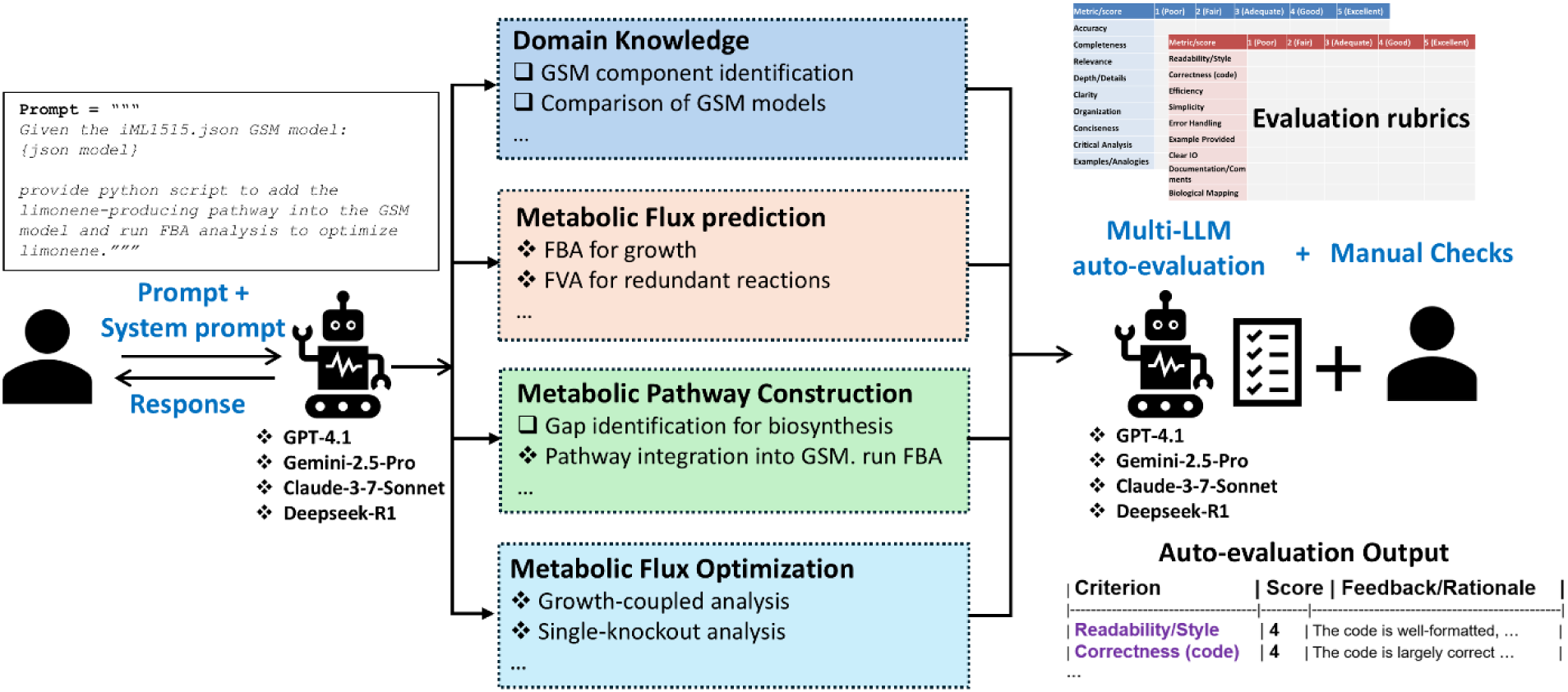

## 1. Introduction

Genome-scale metabolic models (GSMs) have become a cornerstone of modern systems metabolic engineering, providing a systems-level framework for the rational design of microbial cell factories^1–6^. By mechanistically representing the entire network of metabolic reactions and their governing gene-protein-reaction associations within an organism, GSMs enable constraint-based analyses such as flux balance analysis (FBA), which utilizes the stoichiometric matrix and an objective function, to predict cellular phenotypes and cell capacities under various genetic and environmental conditions^6–10^. These simulations are instrumental in the Design-Build-Test-Learn (DBTL) cycle, informing critical engineering strategies and contextualizing experimental outcomes. For instance, they allow researchers to identify competing pathways that divert carbon flux^11,12^, pinpoint precursor and cofactor bottlenecks that limit efficiency^13,14^, and predict optimal gene knockout or overexpression targets^15,16^. By evaluating the feasibility of heterologous pathways and clarifying growth-product tradeoffs, GSMs effectively narrow the vast experimental search space, systematically guiding experimental design toward improved yields, titers, and productivities of valuable chemicals, thereby advancing sustainable bioproduction^17^.

Despite their central role and widespread adoption in academia and industry, the practical application of GSMs is frequently constrained by significant technical and knowledge barriers^18,19^. Developing and executing a robust analysis workflow is a non-trivial endeavor, demanding a confluence of deep biochemical knowledge, systems biology, and computational proficiency. Researchers must navigate complex software packages such as the COBRA Toolbox or COBRApy^20,21^, other relevant tools (PyTFA, OptKnock, FastKnock, OptGene etc.,)^22–27^, and computational algorithms (FVSEOF, iBridge)^28,29^, perform rigorous model curation and validation, and interpret abstract numerical outputs to derive explainable biological insights. This steep learning curve creates a bottleneck, often limiting the direct use of these models to computational specialists and potentially alienating the bench scientists who generate the experimental data and carry out the pathway and strain engineering. Consequently, the full potential of GSMs to drive rapid innovation in bioproduction remains partially untapped. In this study, we leveraged large language models to harness the potential of GSMs.

The recent emergence of Large Language Models (LLMs) offers a potential paradigm shift in how scientists interact with complex computational tools and biological data^30–34^. These LLMs have demonstrated remarkable capabilities in understanding natural language, generating high-quality code, and synthesizing information from vast, unstructured text corpora^30–33,35,36^. In the context of metabolic engineering, LLMs could serve as intelligent “co-pilots”, interpreting the comprehensive network details encapsulated in GSMs, translating high-level scientific questions into executable simulation scripts, explaining the biochemical rationale behind model predictions, and summarizing relevant findings from scientific literature^30,36,37^. Such an interface could dramatically lower the barrier to entry and enhance efficiency, empowering experts and non-experts to directly query GSMs, troubleshoot analyses, and accelerate the cycle of hypothesis generation and testing, thereby democratizing access to these powerful predictive tools.

However, despite this immense promise, there is a lack of empirical evidence regarding the domain-specific competence and reliability of LLMs for the nuanced tasks required in metabolic engineering. To address this critical gap, we present a comprehensive evaluation of LLMs’ capabilities in interpretating GSMs and implementing various GSM related analysis tasks. We systematically benchmarked four prominent LLMs (GPT-4, Gemini, Claude, and DeepSeek-R1) across four main areas: (i) foundational domain knowledge, (ii) metabolic flux prediction, (iii) metabolic pathway construction, and (iv) metabolic flux optimization. Using standardized rubric-based scoring metrics with independent evaluations via a multi-LLM auto-evaluation approach, we identified recurrent failure modes, task- and model-specific capabilities and limitations. Across tasks, the LLMs perform strongly in conceptual explanation and code synthesis, but exhibit consequential failure, including context-window limitations (most pronounced for DeepSeek-R1 and Claude), incorrect assumptions about component identifiers, strain-specific reasoning errors, and errors in domain-specific algorithms. Overall, DeepSeek-R1 performed best on domain tasks, narrowly ahead of GPT-4, Claude, and Gemini. By contrast, the ranking flips for coding: Gemini tops performance, distinguished by robust error handling, clearer documentation, and more readable code, ahead of GPT-4, Claude, and DeepSeek-R1. This work establishes a crucial, evidence-based baseline for LLM-enabled GSM analysis, articulating best practices for current deployment and informing the future development of more reliable, accessible, and automation-ready computational workflow for pathway and strain design.

## 2. Methods

### 2.1. Model selection

We benchmarked four state-of-the-art large language models (LLMs) to better capture diversity in performance across leading Artificial Intelligence (AI) model providers and reasoning paradigms: OpenAI’s GPT-4.1, Google’s Gemini 2.5 Pro, Anthropic’s Claude 3.7 Sonnet, and DeepSeek AI’s DeepSeek-R1. GPT-4.1 is a general-purpose, multimodal model widely adopted for code generation and analytical tasks, featuring a large context window length of 1 million token, and low latency without a reasoning step^38^. Gemini 2.5 Pro is Google’s flagship multimodal model optimized for long-context reasoning and information comprehension, suited to complex document parsing and stepwise analysis, with maximum input token limit of 1 million^39^. Claude 3.7 Sonnet is optimized for strong reasoning, long-context understanding, and reliable instruction following, supporting input contexts up to 200k tokens^40,41^. Deepseek-R1 is a reasoning-centric model optimized for chain-of-thought style reasoning and code synthesis, with a maximum context window length of 128k tokens^42^.

### 2.2. Task Design and Evaluation Scope

We designed distinct tasks to probe LLM capabilities across four key areas within genome-scale metabolic modeling, encompassing domain knowledge, metabolic flux prediction, metabolic pathway construction, and metabolic flux optimization. Domain knowledge covers foundational analyses of GSMs and evaluates LLM fundamental understanding of the GSM’s underlying concepts: summarizing structure; identifying components and subsystems; clarifying biomass composition and growth-coupling concepts; predicting pathways with host selection and bottlenecks; comparing GSMs (including JSON representations); and discussing potential applications, limitations, and potential improvements. Metabolic flux prediction involves applying constraint-based modelling to quantify cellular behavior, using flux balance analysis (FBA) for growth prediction and flux variability analysis (FVA) to characterize redundancy.

Metabolic pathway construction focuses on creating and validating biosynthetic routes: identifying pathway gaps, integrating candidate pathways into GSMs and running FBA, and comparing GSMs for gaps and strain selection. Metabolic flux optimization targets strain engineering strategies, including growth-coupled analyses, flux variability scanning based on enforced objective flux (FVSEOF) to infer up- and down-regulation targets, and single-gene knockout screens and knockout studies for product optimization. Altogether, the study includes 19 distinct tasks executed across four models, yielding 76 in silico experimental runs. To demonstrate real-world utility, we additionally performed nine validation case studies adapted from recent work evaluating microbial cell factories with GSM^6^, expanding the setup to 28 distinct tasks and 112 total in silico experimental runs.

### 2.3. Prompt Design and Model Interaction Pipeline

We designed prompts to yield consistent, automatable outputs across distinct tasks and models using a two-layer prompt architecture: a standardized system prompt message to enforce global behavior and task-specific instructions with explicit descriptions on expected outputs for comparable responses across models to facilitate benchmarking. To parse the GSM into the context, the SBML XML files (retrieved from BiGG model database^43,44^) were first converted into JSON format using COBRApy package^21^ and embedded in the user prompt for interpretation. JSON’s concise key-value and array structure maps naturally to GSM entities (i.e., compartments, genes, metabolites, reactions, stoichiometries, bounds, gene-reaction rules, etc.), eliminating verbose XML tags and attributes for token efficiency and smaller payloads.

A standardized system prompt was employed to anchor model behavior across tasks and LLMs, establishing role, scope, and output constraints before any task-specific instructions. For domain-focused tasks, the system prompt was defined as “*You are an assistant who provides clear and concise response to users*” to prevent verbosity. Whereas for coding-specific tasks, we defined the system prompt as “*You are an assistant who provides clear and concise response to users. Provide full python code first, then only followed by step-by-step explanations. When providing python code, starting with ‘’’python and end with ‘’’. Please think properly to ensure that the code can be run properly with high efficiency*”. This coding-oriented prompt enforces strict formatting and ordering to enable automatic code extraction into python file for downstream execution and analysis. A list of prompts and system prompts used for individual tasks is provided in Supplementary Table S13.

All tasks and models were accessed through NUS AI-Know and interacted via REST API invoked from Python^45^. We set the generation temperature to 0.2 to reduce output randomness and increase determinism and conservatism, which is often desirable in metabolic engineering workflows where scientific correctness, schema compliance, and cross-run reproducibility are critical.

### 2.4. Rubric-based Scoring with LLM Ensemble Auto-Marking and Human Validation

We evaluated the performance of the LLMs on interpreting GSMs and performing analysis tasks using two rubric-based metrics: domain and computational metrics. The domain metric gauges conceptual understanding and scientific knowledge communication across nine criteria: accuracy, completeness, relevance, depth/details, clarity, organization, conciseness, critical analysis, and use of examples/analogies. Each criterion is scored on a 1 (poor), 2 (fair), 3 (adequate), 4 (good), and 5 (excellent/outstanding) scale with explicit rubrics to standardize judgments and ensure comparability across tasks and models (Supplementary Table S1). The computational/coding metric assesses code-centric outputs across nine criteria: readability/style, correctness (code), efficiency, simplicity, error handling, example provided, clear IO, documentation/comments, and biological mapping. Scores are likewise assigned on a 1-5 scale using detailed rubrics (Supplementary Table S2).

To improve auditability, we employ an ensemble of the four LLMs to automate the marking process independently following the same rubrics embedded in the prompt. Based on a single task-based instruction prompt and the given response from the individual LLM, each LLM provides score of 1-5 for each criterion supported with rationale in a tabular format. The prompt template is provided in Supplementary Figure S1.

Beyond LLM auto-marking, we performed human, end-to-end validation for all tasks to ensure factual accuracy and correct interpretation of GSM content. For coding analysis tasks, we executed the generated code extracted from the response in a controlled environment with the required packages installed to verify if the code can run properly, adheres to specified inputs and analysis sequences, and produces the expected results. Failures or deviations were traced to identify the specific implementation issues. We also audited LLM’s scores and per-criterion rationales, verifying that each justification accurately reflects the response and task requirements.

Crucially, this human-in-the-loop audit served as a calibration mechanism to identify structural limitations inherent to multi-LLM auto-assessment. Because the evaluator models share overlapping training distributions, the ensemble remains bounded by common data blind spots. To systematically isolate these occurrences, we tracked instances where the intra-ensemble scoring variance approached zero (s^2^ ➔ 0), but human verification revealed operational or semantic failures. These instances were classified as “Consensus Hallucinations”, cases where the automated judges universally misjudged or overlooked hidden faults. This cross-validation framework proved essential for tracking the degradation of LLM auto-marking reliability when transitioning from heavily documented model organisms to less pervasive datasets.

## 3. Results

### 3.1. Benchmarking workflow

Figure 1 illustrates the benchmarking workflow developed to evaluate LLMs’ capabilities in interpreting and analyzing GSMs. We benchmarked four leading LLMs: OpenAI GPT-4.1, Google Gemini 2.5 Pro, Anthropic Claude 3.7 Sonnet, and DeepSeek-R1 to capture performance diversity across reasoning paradigms. We evaluated the LLMs across four capability areas: (i) domain knowledge, (ii) metabolic flux prediction, (iii) metabolic pathway construction, and (iv) metabolic flux optimization. The domain knowledge tasks assess foundational GSM understanding including model structure summarization; identification of components and subsystems; explanation of biomass composition and growth-coupling; pathway prediction with host selection and bottleneck analysis; comparison of GSMs; and discussion of applications, limitations, and potential improvements. Flux prediction tasks involve implementing constraint-based modelling, using flux balance analysis (FBA)^7,9^ for growth prediction and flux variability analysis (FVA)^46^ to characterize redundancy. Pathway construction tasks address gap identification for a specific pathway, integration of candidate biosynthesis pathway into GSM followed by FBA validation, and model comparison to identify gaps and guide strain selection. Flux optimization tasks target strain engineering via growth-coupled analysis, flux variability scanning based on enforced objective flux (FVSEOF)^29^ to infer up- and down-regulation targets, and gene knockout screening for product optimization. In total, 19 distinct tasks were executed by the four LLMs respectively, yielding 76 in silico experimental runs.

**Figure 1:**
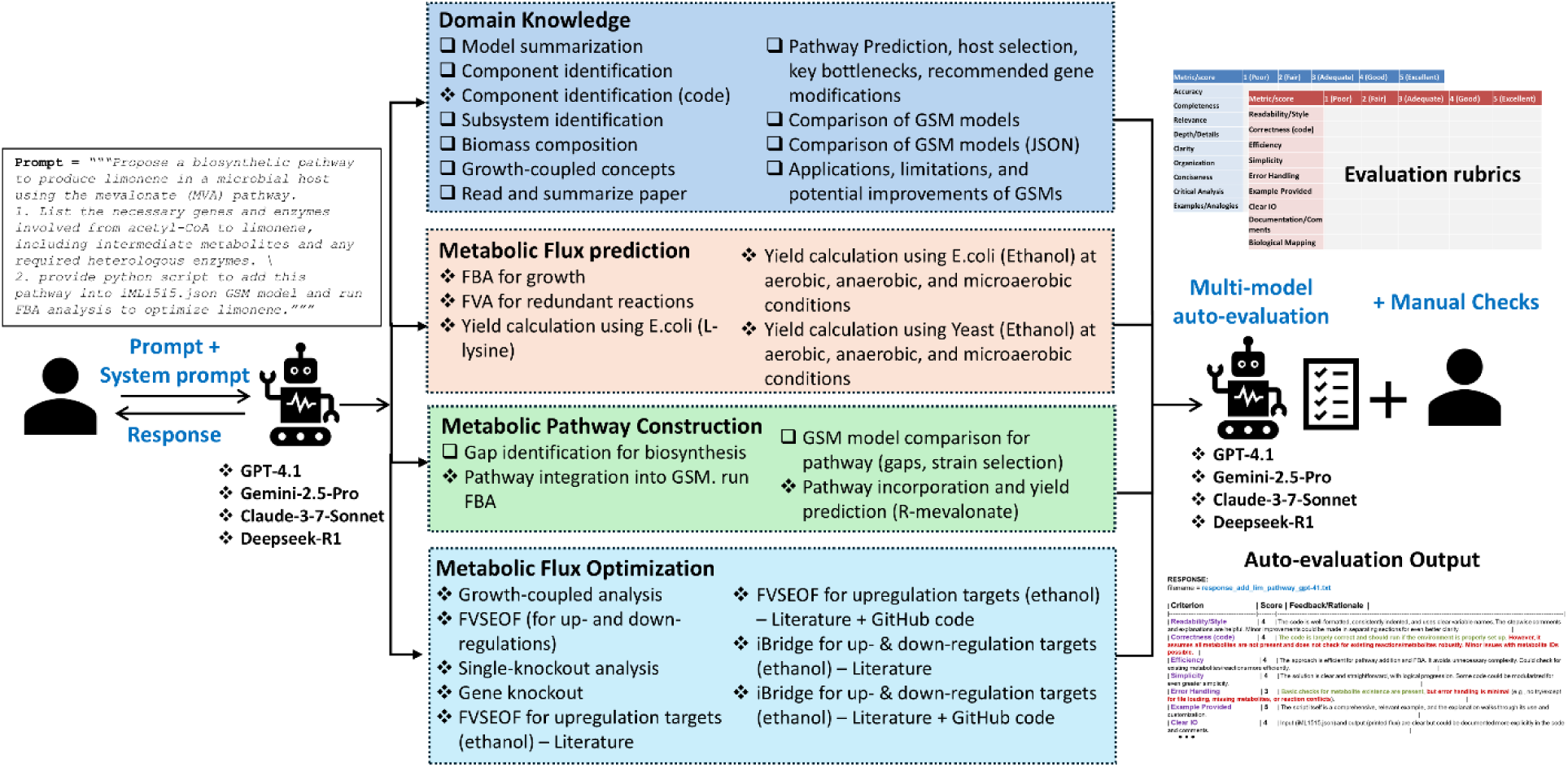
Overview of the benchmarking workflow: An instruction prompt and a standardized system prompt for overarching instruction (role, scope, formatting) were executed across four prominent LLMs (GPT-4.1, Gemini-2.5-Pro, Claude-3.7-Sonnet, DeepSeek-R1) on tasks designed spanning four core areas in interpreting and analyzing GSMs. The prompt and response were then auto-evaluated using rubric-based criteria by all four LLMs independently, with human-in-the-loop verification. The square bullets represent domain tasks, while the diamond bullets indicate coding tasks.

We evaluated the LLMs’ performance on interpreting GSMs and performing analysis tasks using two rubric-based metrics: (i) a domain metric for conceptual understanding and scientific communication (accuracy, completeness, relevance, depth, clarify, organization, conciseness, critical analysis, and use of examples/analogies); and (ii) a computational/coding metric for code-focused outputs (readability/style, correctness, efficiency, simplicity, error handling, examples, clear I/O, documentation/comments, and biological mapping. Each criterion was scored 1-5 with explicit rubrics to standardize judgements (Supplementary Table S1-S2). To improve auditability, we used an ensemble of the four LLMs to auto-mark the prompt and response independently against the same rubrics, with tabulated per-criterion scores and rationales. Beyond auto-marking, we conducted human end-to-end validation: verifying factual accuracy and GSM interpretation; executing extracted analysis code in a controlled environment to confirm compliance with inputs, required analysis steps, and expected outputs; tracing any failures to specific implementation issues; and auditing LLM-assigned scores and rationales to ensure accurate reflection of the task and responses. In the following sections, we elaborate the findings across major areas (domain knowledge, metabolic flux prediction, and others).

### 3.2. Domain Knowledge

For domain knowledge tasks, we assess the fundamental knowledge of LLM on the factual concept of genome-scale models/modelling, and the reasoning capability in interpreting genome-scale model input files. For interpretation-related tasks, we converted the SBML XML file of the GSM into the JSON format and embedded it into the prompt for LLMs. The first task involves summarizing the details of the model and providing the model structure in a visual tree. Gemini (39.25) covers the major sections (metabolites, reactions, genes, compartments, and GSM organism) with counts and key information and illustrates the hierarchical organization of the JSON data with effective use of examples. Instead of providing the actual outputs, DeepSeek (40.75) and GPT (35.5) provide the python code to extract the details from the model (ID, version, compartments, gene count, reactions, etc.) and parse the data structure correctly. These LLMs provide a functional approach to summarize and visualize a GSM but lack deeper explanations into the model components and insights into their biological significance, and limited critical analysis of the model structure, approach, and specific aspects to highlight. Claude is not applicable for this case with GSM input due to context window limit.

For the component identification task, LLMs were prompted to identify reactions involving a specific compound as substrate and summarize the involved pathway. There are major gaps in the responses for LLMs: (i) list of reactions is incomplete with notably missing significant reactions; (ii) inaccurate with the compound as product or the compound is not involved at all; (iii) pathway could be wrong due to the earlier errors; (iv) lack evaluation of metabolic role and significance. For this task, all LLMs have weak to moderate performance with an average mean score of 36.33 (GPT: 37.75; DeepSeek: 36.50; Gemini: 34.75). Again, Claude is not feasible for this task with GSM input limited by context window constraint.

To address the concern on accuracy, we assessed the LLMs’ ability to perform the component identification task using code-based approaches, employing same prompt but with a modified system prompt that enabled python code generation. Only GPT produced functional code that generated accurate results, including proper JSON file loading, identification of reactions involving acetyl-CoA as a substrate (represented with negative coefficients), collection of reaction details, tabular display of results, pathway (subsystems) summarization based on reaction counts, and comprehensive step-by-step explanations. This led to a higher score of 38 for GPT, with points deducted for limited error handling (Figure 2c-d), such as error check for missing file or missing mandatory field keys. DeepSeek’s implementation contained critical errors, including incomplete code for file loading and incorrect assumptions of EcoCyc^47^ as pathway identifier without validating other pathway annotations. Despite these issues, DeepSeek (39) scored well on efficiency, example provided, clear IO, and biological mapping. Gemini (36.25) performed weakest in this evaluation, providing truncated JSON input strings and limited examples, resulting in lower performance across most criteria (Figure 2c).

**Figure 2:**
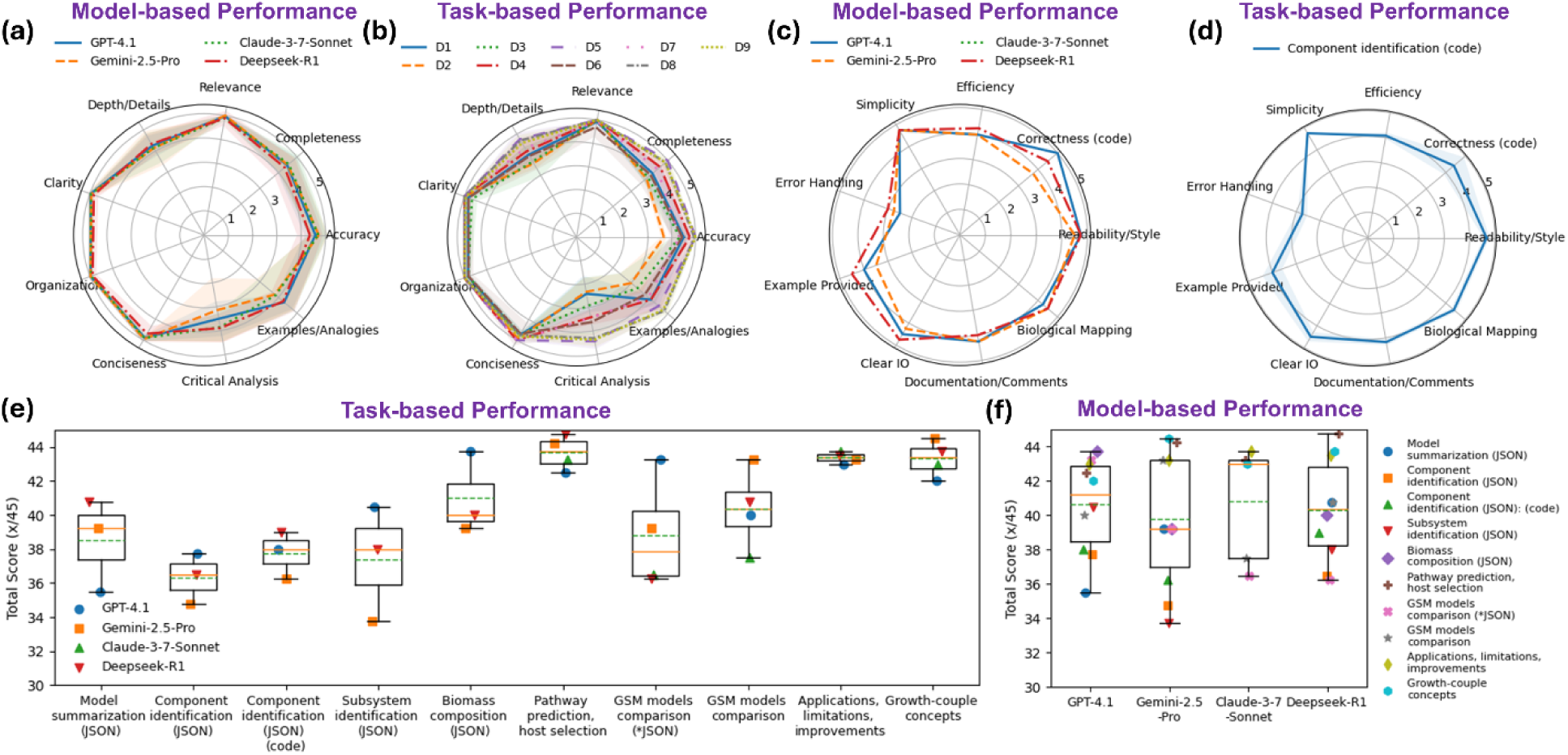
Evaluation of domain knowledge tasks (a) Model-based performance across evaluation criteria of domain tasks. (b) Task-based performance across evaluation criteria of domain tasks. (c) Model-based performance across criteria of coding tasks. (d) Task-based performance across criteria of coding tasks. (e) Consolidated score for task-based performance. (f) Consolidated score for model-based performance. D1-D9 denotes the specific domain tasks labelled in (e) from left to right, excluding component identification (JSON) (code).

The LLMs were also assessed in their capabilities to read the GSM and summarize the reactions or pathways for a subsystem (e.g., cofactor NAD+/NADH or NADP+/NADPH). While covering broad range of reactions and pathways with good depth and roles explanation, most LLMs’ responses (GPT (40.5) and DeepSeek (38)) contain significant error of omission for some key reactions and contain some inaccuracies and inference without direct evidence. Gemini (33.75) provides only factual listing, lacks details of reaction function, pathway aspect of the system, and analysis and explanations of the roles in the metabolic context. Again, Claude is not applicable for this case due to limited window capacity.

Biomass is often the primary or constrained objective for GSM flux analysis to predict the maximum possible growth rate under given conditions^48^. We thus assessed the LLMs’ capabilities in explaining the composition and role of the biomass objective function in the GSM model, and the key differences between the two biomass functions (WT and core biomass). In general, all LLMs, except Claude (due to context window limit), accurately explained the role, composition, and differences between the biomass functions (detail level, components included and use case). GPT (43.75) scored the best with full mean scores for most individual criteria except critical analysis; addressing all aspects more thoroughly, including appropriate implementation technical details, examples of the stoichiometry coefficients and a comparative summary table that effectively illustrates the differences which are lacking in other models (DeepSeek: 40, Gemini: 39.25). The responses could be improved with explicit critical evaluation of limitations (e.g., static nature, condition-dependency).

Given the role of GSMs in pathway and strain engineering, it is essential to evaluate LLMs’ capabilities in proposing comprehensive biosynthetic pathway designs. To this end, the assessment focused on limonene production via mevalonate pathway (as a case study)^49^ requiring the model to (i) list the necessary genes and enzymes involved, (ii) suggest possible host, (iii) highlight key regulatory points and potential bottlenecks, and (iv) recommend gene modifications to enhance yield. The results show that all the LLMs correctly described the mevalonate (MVA) pathway with accurate genes, enzymes, and intermediates^49^. The models appropriately suggested *Escherichia coli* due to its rapid growth and genetic tractability, *Saccharomyces cerevisiae* with native MVA pathway, and *Yarrowia lipolytica* for high acetyl-CoA flux, align with established literature^50–52^. The responses also demonstrated sound critical evaluation in identifying the bottlenecks, including acetyl-CoA supply limitations, HMG-CoA reductase (HMGR) as the primary rate-limiting enzyme due to feedback inhibition, precursor competition at the GPP branch point, IPP/DMAPP balance requirements for GPP synthesis, limonene toxicity, and cofactor imbalance from NADPH requirements for HMGR^53–55^. Correspondingly, the models proposed potential targeted solutions including (i) enhancing acetyl-CoA supply, (ii) overexpressing bottleneck enzymes (using truncated and deregulated HMGR, optimizing GPPS/LS expression with codon-optimized genes and strong promoters), (iii) knocking down FPP synthase to reduce GPP consumption, and (iv) mitigating limonene toxicity via efflux pump or two-phase fermentation with organic overlay for separation^56^. All models performed well on this task, with consolidated scores >= 42.5. DeepSeek achieved near perfect performance (44.75/45), providing concrete example strain designs for both *E. coli* and *S. cerevisiae* that were well-integrated in the response, followed closely by Gemini (44.25), Claude (43.25), and GPT (42.5).

Strain selection requires comparative analysis across multiple GSMs to identify optimal performers. To evaluate this capability, LLMs were evaluated on their ability to read and compare key metabolic reactions/pathways differences between two GSMs: iML1515^57^ for *E. coli* and yeast850^58^ for *S. cerevisiae*. Due to the context window constraints, only reaction name, subsystem (pathway), and metabolites stoichiometric data were parsed from the two GSMs. GPT demonstrated superior performance in this comparative analysis task, successfully reading and interpreting information from both models, covering all major and minor metabolic pathway differences including unique features, and providing a well-structured summary table delineating distinctions coupled with appropriate concluding remarks, earning GPT the highest score (43.25) across all evaluation criteria. The response could be enhanced with more functional implications of the identified differences, specific reaction details and relevant analogies. While Gemini (39.25) successfully read both GSMs, its response was more limited in scope, covering only some major pathways with predominantly descriptive content, lacking coverage of minor pathways, detailed reaction/pathway, critical analysis into model strengths and limitations, and comparative analogies. In contrast, both Claude (36.5) and DeepSeek (36.25) encountered significant limitations, only able to interpret one GSM model, thus failing to provide essential comparative analysis between the two GSMs.

To address context window limitations for most LLMs, we evaluated their fundamental knowledge by comparing two GSMs’ details and key metabolic reactions/pathways differences without providing the actual GSM input data. While all models successfully presented model statistics in tabulated format, and key pathway differences, their responses contained minor but significant deviations in the quantitative data (gene, metabolite, reaction counts) particularly for yeast850 (Supplementary Table S10). The qualitative pathway comparisons were mostly accurate but lacked deep explanations of their biological implications, illuminating analogies, and critical analysis of model strengths and limitations. Gemini distinguished itself in this task by providing good analysis of the implications of model differences and practical applications for engineering tasks in its concluding remarks, achieving the highest score (43.25). In contrast, Claude scored the lowest across completeness, depth/details, critical analysis and use of examples/analogies criteria, yielding the lowest overall score of 37.5, behind DeepSeek (40.75) and GPT (40)

Growth-coupling bioproduction (GCBP) is a well-established metabolic engineering strategy that creates an obligatory dependency between product formation and essential cellular metabolic functions, ensuring a high minimal yield and genetic stability^59–61^. We thus assessed LLMs’ knowledge of growth-coupling engineering concepts across five dimensions: (i) definition: explaining what it means to be growth-coupled and contrast strong and weak growth coupling; (ii) analysis: describing how GSM can be used to determine if a pathway is growth-coupled, and suggesting and comparing tools/methods; (iii) engineering relevance: why is growth-coupling important in metabolic/strain engineering and support with successful examples; (iv) limitations and challenges: what are limitations of using in silico growth-coupling analysis and how can experiment observations diverge from model predictions; (v) future directions: opportunities for improving growth-coupled strategies computationally and experimentally.

All LLMs demonstrated expert-level understanding of growth-coupling in metabolic engineering, providing comprehensive and technically precise explanations that addressed all aspects of the prompt with appropriate depth and clarity. The models exhibited strong knowledge of the key analytical and design methods: production envelope (phenotypic phase plane)^62,63^, FSEOF (flux scanning based on enforced objective flux) for analyzing given designs, and bi-level optimization (OptKnock, RobustKnock, OptForce)^25,26,64^ for identifying optimal designs. They demonstrated familiarity with specific tools including the COBRA toolbox (FBA, FVA) for analysis, and specialized but computationally intensive design tools such as OptKnock, OptCouple, gcOpt, and FastPros^65–68^. Highly effective examples (succinic acid in *E. coli*, lactic acid in yeast, lysine in corynebacterium glutamicum, 1, 4-Butanediol (BDO) in *E. coli*) were integrated seamlessly in the explanations. The models demonstrated deep critical thinking in proposing future directions, including multi-omics integration for model validation and refinement, machine learning applications for predicting and optimizing growth-coupling designs, CRISPR-based genome editing for precise multiplexed modifications, genetically encoded biosensors for high-throughput screening, and adaptive laboratory evolution (ALE) for strain optimization. Gemini achieved the highest score (44.5/45), followed by DeepSeek (43.75/45), Claude (43/45), and GPT (42/45), with minor improvements suggested for latter models by including more algorithmic differences, tool-specific details, critical trade-off analysis between approaches, and deeper mechanistic insights.

Lastly, LLMs were assessed for their fundamental knowledge and reasoning capabilities to provide detailed analysis of the role of GSM covering the applications on “How” with successful examples, limitations by highlighting the key challenges and how this affects prediction accuracy, and improvements by discussing opportunities for improving predictive power and future directions. All LLMs responded well to the prompt, providing in-depth information supported by highly relevant examples and references, along with strong critical analysis and insightful discussion. As a result, consistently high consolidated mean scores of >= 43 out of total 45 marks were achieved. The models demonstrated thorough understanding of GSM applications, including predicting metabolic flux distributions and theoretical yield, evaluating effects of gene modifications such as gene knockouts, overexpressions, and insertions, guiding metabolic engineering designs with desired phenotypes, utilizing OptKnock for identifying gene deletion strategies that couple growth with production, identifying bottlenecks and optimal intervention points for pathway enhancement, balancing cofactors, and selecting most promising host. The responses were supported with concrete examples: including the engineering of *E. coli* for succinate production using GSM-guided knockouts^26^, optimization of mevalonate pathway for enhanced isoprenoid production in *S. cerevisiae*, balancing precursor supply and minimizing byproducts of 1,4-butanediol in *E. coli* etc.

LLMs have identified comprehensive limitations in GSM, including structural, data-quality issues, and computational challenges. Key challenges include incomplete genome annotation, inaccurate network reconstructions, lacking regulatory and kinetic information, and reliance on steady-state assumption. Additionally, assumptions in objective functions such as maximizing growth or specific metabolite production may not reflect the true cellular priorities. Computational challenges further complicate model performance including vast solution space that hinder unique predictions, parameter uncertainty (kinetic parameters are often unavailable, unconstraint lower and upper bounds), and integration challenges with omics data. These limitations can contribute to discrepancies between computational predictions and experimental outcomes. Several approaches were proposed by LLMs as potential solutions to these challenges: Integrating multi-omics data for better model constraints; regulatory/kinetic modelling (metabolism and expression ME-models (COBRAme)^69^, GECKO (include enzymatic constraints)^70^, dynamic and whole-cell models); machine learning techniques to fill knowledge gaps through sequence to function prediction for model reconstruction and learning complex regulatory rules from omics data; automated model reconstruction tools (CarveMe, ModelSEED)^71–73^; and community modelling, hold promise for enhancing model accuracy and expanding GSM utility in metabolic engineering. A table summarizing the key insights derived from different LLMs is available in Supplementary Table S11.

Across domain-knowledge tasks, all LLMs performed strongly, achieving consolidated mean scores above 4 for most criteria. The main weak spots were critical analysis and examples/analogies, which also showed greater variability (Figure 2a-b). Overall performance was consistently high: 70% of tasks (7 of 10) scored ≥ 38.5 (85% of total marks), reflecting a robust grasp of core GSM concepts (e.g., biomass objectives, pathway design, host selection, growth-coupling, and GSM applications and limitations), with weak performance in component and subsystem identification (Figure 2e). The main limitation lies in GSM-input reasoning reliability. Models struggled to interpret large GSM files and reliably extract details, frequently omitting key reactions/pathways, making unverified assumptions about identifiers, and inferring biological interpretations without direct evidence from the provided data. Careful validations via code-based extraction and expert review remain essential. Among models, Claude achieved the highest mean score (40.8), narrowly ahead of GPT (40.625), DeepSeek (40.325), and Gemini (39.775) (Figure 2f). However, Claude failed five GSM-input tasks due to context-window constraints, limiting its practical robustness. A tabulated summary of key insights derived across all models for all the domain-focused tasks is provided in Supplementary Table S3.

### 3.3. Metabolic Flux Prediction

Flux Balance Analysis (FBA) and Flux Variability Analysis (FVA) are core methods within constraint-based modelling of genome-scale metabolism, where kinetic parameters are largely unknown^9,46^. FBA is applied to GSMs that assumes steady-state and uses stoichiometric and capacity constraints to compute a flux distribution that optimizes a specified objective (e.g., biomass growth or metabolite production). It yields one optimal solution, but often there are many alternative optima with the same objective value. Beyond just a single optimal solution found by FBA, FVA systematically explores the solution space by calculating the minimum and maximum feasible flux for each reaction while still satisfying overall network constraints.

We evaluated the LLMs capabilities in generating python code to run FBA analysis on GSM (e.g., iML1515.json) with explanations and identify top 10 reactions with the highest flux. All models score highly for correctness; they use the COBRApy library appropriately to load the model^21^, execute FBA, print the objective value (e.g., biomass flux), sort and print the top reactions with relevant details. The readability/style and simplicity are also strong, where the code and explanations are clear, well-formatted, and easy to follow, while the solution from the code implementation is elegant and minimal yet being comprehensive. The greatest variability among the responses by LLMs is in error handling: Gemini performed the best (42.25) with robust try/except handling for missing file, solution status, and general exceptions with helpful error messages and instructions, whereas GPT and Claude omitted these safeguards, leaving the code more fragile for real-world applications. Manual inspections confirmed that all the codes were able to run FBA without errors and produce correct results.

We further assessed LLMs on generating code to run FVA on a GSM model with explanations and identify the top 10 redundant reactions with the largest flux range. These reactions typically indicate alternative pathways or cycles in the metabolic network that can carry different flux values with high metabolic flexibility without significantly impacting the objective function (growth rate). All codes were implemented using appropriate functions from the COBRApy package, performing FVA with 90% or 100% of optimal growth with expected results without errors. LLMs have comparable performance across criteria, though lower mean error handling scores with no or minimal error handling (i.e., only checks for the model file) for most LLMs. The biological relevance was well-addressed by correctly mapping large flux range to the concept of metabolic redundancy and providing reaction names for enhanced interpretation.

In summary, both FBA and FVA tasks achieved comparable mean scores of roughly 39/45 (Figure 3c). Gemini is the top performer in both (FBA: 42.25, FVA: 40.25), particularly excelling in error handling and documentation/comments (Figure 3a). This is followed by DeepSeek, tied for second across tasks, scoring 39.75 for FBA and 39.25 for FVA task with minimal error handling. While Claude and GPT attained lower mean scores for both tasks, they remained reliable in correctness, simplicity and efficiency in implementing the tasks. Overall, the LLMs delivered a well-structured, correct implementation of FBA and FVA analyses using COBRApy^21^, with efficient code and appropriate use of pandas for data manipulation and sorting. The generated code was exceptionally readable with clear comments and comprehensive step-by-step explanations that bridge computational steps and biological significance (Figure 3b). Despite no explicit example output is shown, the code structure makes expected output evident. The main weakness is limited error handling (Figure 3b) which could be improved by adding more robust error handling for file operations and optimization failures. A tabulated summary of performance evaluation across models and tasks is available on Supplementary Table S4.

**Figure 3:**
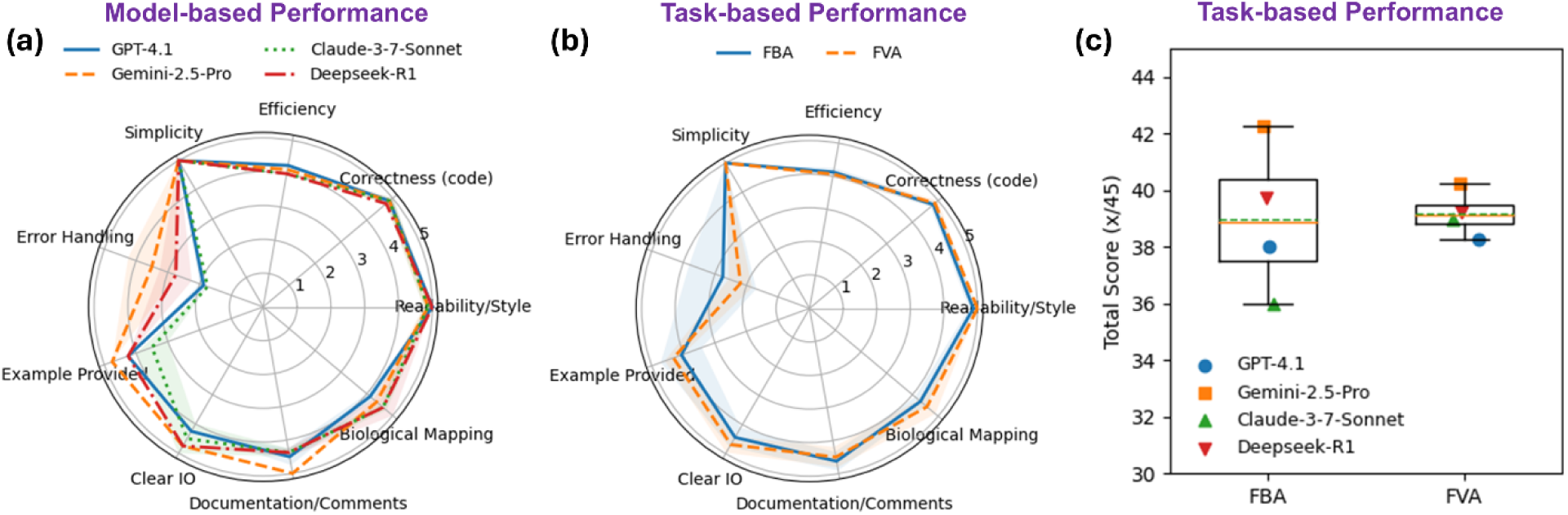
Evaluation of metabolic flux prediction tasks (a) Model-based performance across varied criteria for coding tasks (b) Task-based performance across evaluation criteria for coding tasks. (c) Consolidated score for the four models across tasks.

### 3.4. Metabolic Pathway Construction

To evaluate performance on pathway construction, we first assessed the LLMs’ ability to identify gaps in the GSM for a specific pathway. LLMs were prompted to read the GSM JSON file and suggest candidate reactions and enzymes that could fill metabolic gaps to allow for compound limonene synthesis. DeepSeek delivered the most comprehensive response, addressing all aspects: gaps, candidate reactions, heterologous enzymes, additional considerations (e.g., enhancing precursor supply, localization, model updates), and providing a sample pathway addition; achieving the highest scores across most criteria and the highest total score (43.75/45). However, the model discussed only the methylerythritol phosphate (MEP) pathway without considering alternatives such as mevalonate (MVA)^56,74^. Similarly, Gemini focused solely on MEP: identified the missing gap and suggested the required enzymes, but offered minimal details on candidate reactions, enzymes/EC numbers, and lacked deeper evaluation of the proposed solution, alternatives, implementation considerations, and analogies, resulting in a lower score of 37.25. In contrast, GPT’s response covered heterologous reactions to native MEP (with enzymes and source organisms), the alternative MVA pathway, and provided a sound analysis of upstream precursor limitations and competing pathways like FPP synthesis, yielding strong overall performance (41.75; median 43.5). Limited by the context window constraint, Claude is not applicable for this task with GSM JSON input.

Strain comparison and selection are important in metabolic engineering for optimizing production yield^6^. To evaluate this, we prompted LLMs to compare two GSMs (iML1515 and yeast850^58^) for limonene synthesis by identifying the metabolic gaps, determining which strain is better, and proposing strategies to improve the yield. To avoid infeasibility due to context-window constraints, we relied on the models’ fundamental knowledge rather than explicitly embedding the GSM files. All LLMs addressed these three aspects thoroughly: (i) identified the gaps after considering the native MEP and MVA pathways, (ii) selected yeast as the superior strain due to higher precursor fluxes and tolerance while articulating the pros and cons for each strain, and (iii) proposed yield-improvement strategies such as enhancing precursor supply, blocking competing pathways, and cofactor regeneration etc. (refer Supplementary Table S12 for details), resulting in a high average score of 42.15. Again, DeepSeek delivered outstanding performance with more in-depth technical analysis (e.g., compartmentalization, cofactor imbalances), descriptions of in silico methods (FBA/FVA), effective examples including quantitative data from literature (highest titers for each strain), and specific gene knockouts, yielding the top score of 43.25. This was followed by Gemini (42.5), Claude (41.5), and GPT (41.25), whose lower scores reflect omission in certain listed aspects.

After the two domain-focused tasks (gap identification and strain comparison), we evaluated LLMs’ implementation capability to integrate a pathway into the GSM model. We prompted LLMs to first propose a biosynthetic route for limonene production via the MVA pathway from precursor acetyl-CoA, listing the necessary genes, enzymes, intermediate metabolites, and any required heterologous enzymes; next provide python code to add the pathway to the iML1515 GSM model and run FBA to optimize limonene. All LLMs correctly proposed the right pathway with nine reactions (Figure 4a). However, on the coding performance, only GPT produced fully functional code that added the complete pathway and executed flux analysis properly; the other models generated codes with errors such as incomplete reactions, wrong metabolite ID assumptions, stoichiometric imbalance (e.g., proton), and incomplete codes for pathway incorporation. Overall, performance on pathway integration into GSM was unsatisfactory (Figure 4g), with an average score of 34.88 (median: 37.13), ranked from GPT (38.75), Gemini (38.75), Claude (35.50), to DeepSeek (26.5). Despite being functional, GPT’s code lacked robust checks for existing reactions/metabolites, missing metabolites, reaction conflicts, and file loading, slightly lowering its score. In contrast, Gemini scored higher on error handling, readability, and documentation/comments, which offset some implementation errors, yielding a comparable average score as GPT. DeepSeek’s unexpectedly low score was driven by multiple syntax errors (e.g., incorrect dictionary keys, invalid metabolite definitions), lack of error handling, and incomplete codes, likely attributed to the limited context window and the extended chain-of-thought while solving complex tasks.

**Figure 4:**
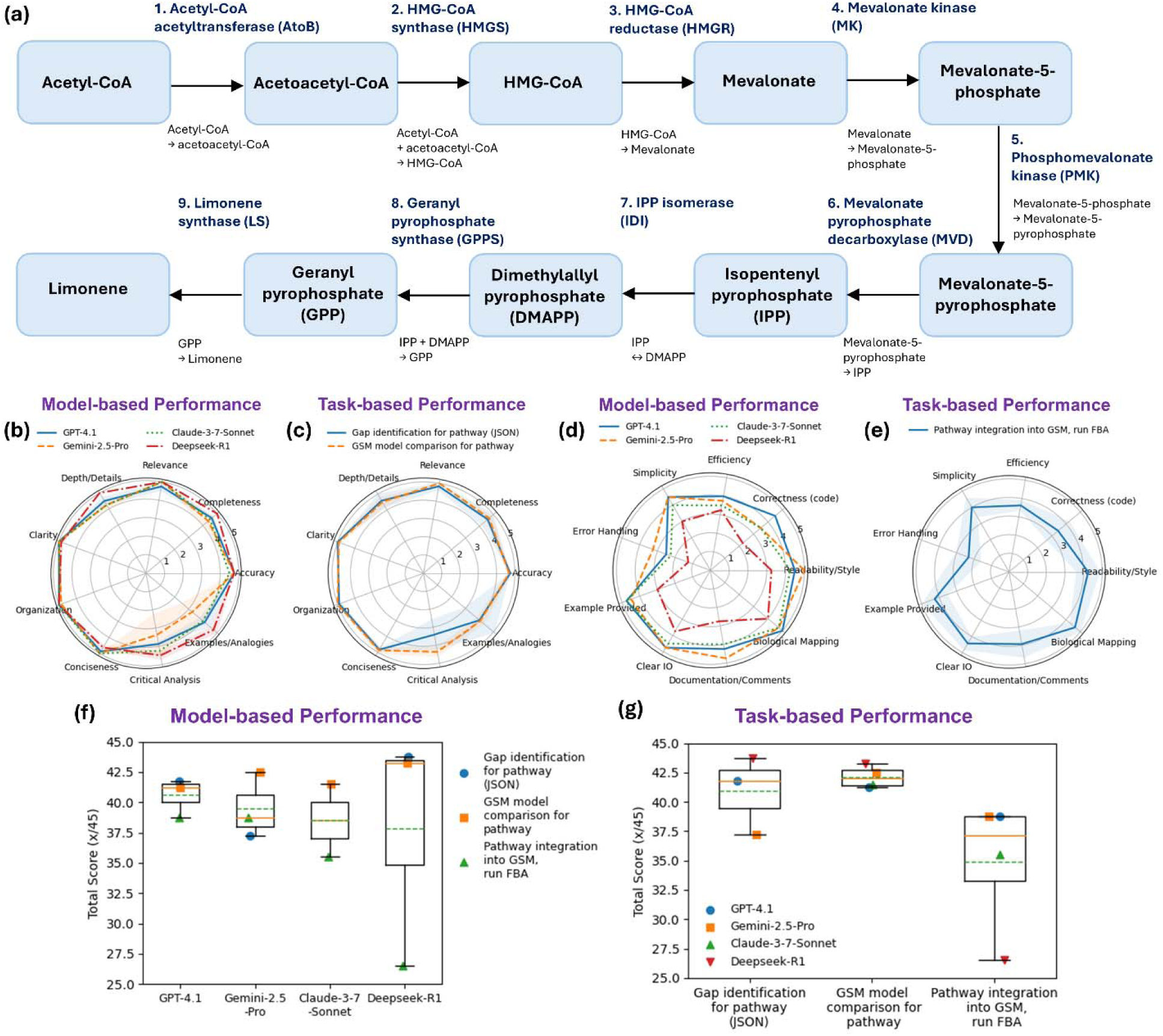
Evaluation of metabolic pathway construction performance (a) Limonene-producing pathway as a case study (b) Model-based performance across evaluation criteria for domain-specific tasks. (c) Task-based performance across evaluation criteria for domain-specific tasks. (d) Model-based performance across different evaluation criteria for coding tasks. (e) Task-based performance across evaluation criteria for coding tasks. (f) Consolidated score for model-based performance. (g) Consolidated score for task-based performance.

To sum up, for domain-specific tasks involving gap identification and GSM model comparison for pathways (without JSON input), DeepSeek led nearly all evaluation criteria (Figure 4b-c) for model-based performance, achieving the highest average score (43.50), followed by Claude (41.50) and GPT (41.50), and Gemini (39.88). The greatest inter-model variability was observed in critical analysis and the use of examples/analogies, followed by depth/details and completeness. For the coding-specific task of integrating pathway into the GSM, all LLMs performed weakly (Figure 4d-e); only GPT produced functional code that was executed without errors, whereas Gemini and DeepSeek scored low on correctness. Across models, insufficient error handling was a recurring limitation, exceptionally critical in this task, where rigorous validation of metabolite and reaction IDs is essential to prevent missing compounds and avoid conflicts with native components. Based on the consolidated average score across model-construction tasks, GPT showed the best overall performance when considering both domain- and coding-specific tasks, followed by Gemini, Claude, and DeepSeek (Figure 4f). For tasks requiring ingestion of GSM JSON input or extended context for complex, multi-step reasoning, Claude and DeepSeek were suboptimal choices due to context window limitation. A table summarizing key insights for task- and model-based metabolic pathway construction category is also provided in Supplementary Table S6.

### 3.5. Metabolic Flux Optimization

Flux Variability Scanning based on Enforced Objective Flux (FVSEOF) is a technique used in metabolic engineering to identify up- or/and down-regulation targets that enhances metabolic flux of a specific target compound^29^. We evaluated LLMs on implementing FVSEOF to identify up/down-regulation targets to improve succinate production in *E. coli* GSM (iML1515). From preliminary analysis, there were significant variabilities in the specific methods used for the algorithm across models. To standardize the method, we defined the following procedure: (1) at each enforced production level, use FVA to obtain min/max fluxes; for each reaction, (2) calculate the correlation between enforced level and the min flux; (3) identify and report top 10 reactions whose min flux is most positively correlated (upregulation targets), and top 10 most negatively correlated reactions (downregulation targets)^6^. Gemini scored highest (41.75) (Figure 5a, d) with correct use of context managers for temporary modification of model, loopless FVA, biomass viability, NaN handling when computing correlation, and strong code comments, though loopless FVA slowed execution. Claude (40.75) also applied context managers, enforced biomass viability, parallelized using the ‘processes’ argument in FVA (4 CPU cores) for efficiency, plotted correlation distribution and flux profiles, and saved results into CSV files. GPT (38.0) was concise with fast operation (use slim_optimize() from COBRApy^21^ when computing max target flux for efficiency), but lacked biomass consideration, relied on full model copies instead of context managers, and had sparse comments. DeepSeek (37.75) failed to isolate constraints across iterations (no copy/context manager) which distorted the results, omitted infeasibility flux check after FVA and produced NaNs for downregulation. Across models, explicit ‘try/except’ checks for file I/O, reaction IDs, solver errors, and NaN correlations could improve reliability.

**Figure 5:**
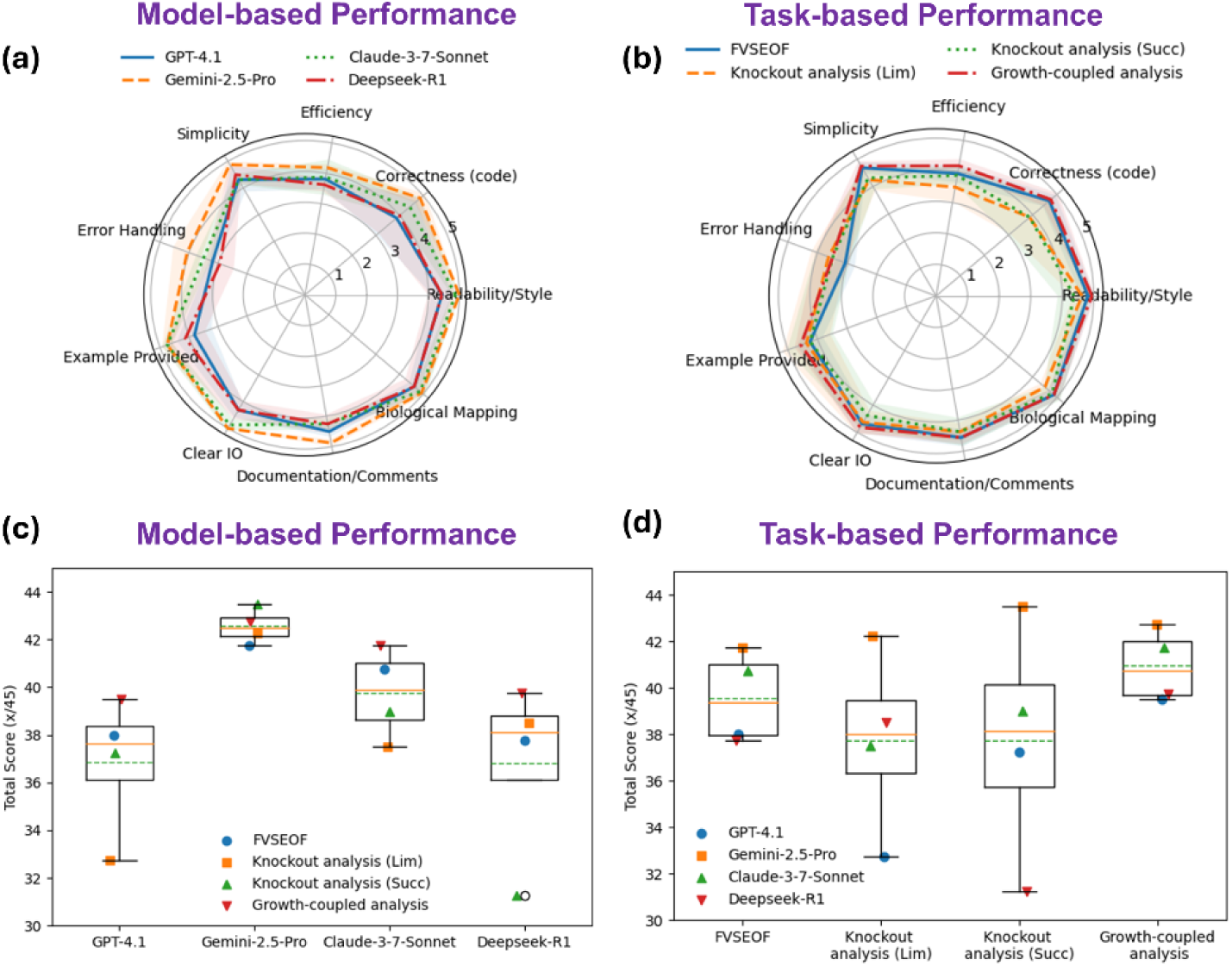
Evaluation of metabolic flux optimization tasks (a) Model-based performance across evaluation criteria for coding tasks (b) Task-based performance across criteria for coding tasks. (c) Consolidated model-based score. (d) Consolidated task-based score.

Knockout analysis is a core technique in GSM for identifying gene/reaction deletions that redirect flux toward a desired target. We assessed LLMs on single-knockout analysis to improve product yield (e.g., limonene) while maintaining 10% of maximal biomass. Again, Gemini delivered production-quality code (42.25) (Figure 5a, c-d), with excellent documentation, informative error messages, and print statements for clarity, parallelized knockout runs, and safe constraint handling via context managers; a minor improvement would be verifying reaction IDs before use. DeepSeek placed second (38.5), incorporating solution optimality check, but relying on inefficient manual loop through genes and omitting wild type comparisons for improvement validation and reactions IDs verification. Claude achieved a comparable score (37.5), validating IDs and solution optimality, performing single gene and reaction deletions via manual loop with a plus feature from its CSV outputs and plots. GPT produced non-functional code (32.75), due to misuse of the built-in ‘cobra.flux_analysis.single_gene_deletion’ function^75^ (wrong key assumptions) and redundant model loads; while it included basic checks for biomass and product reactions, the model lacked infeasibility check for solver optimality.

We further evaluated knockout analysis workflow for a native-pathway target (e.g., succinate production) using the iJR904 E. coli K-12 model, a common testbed for optimizing succinate yields via gene deletions ^76^. LLMs were prompted to (i) suggest genes to knock out to enhance succinate production under anaerobic conditions, (ii) run FBA to identify succinate flux when optimizing biomass anaerobically, (iii) run FBA to optimize succinate while maintaining 10% optimal biomass flux, and (iv) implement the individual suggested gene knockouts using COBRApy to determine the impacts on yield and biomass. Despite following the intended workflow, all LLMs failed in execution with at least an error, due to incorrect biomass reaction ID. Gemini achieved the highest score (43.5) across all evaluation criteria, aided by safe model context managers and robust handling of model loading and solution status check, but it assumed an incorrect biomass ID and lacked validations in genes IDs. Claude ranked second (39.0) but exhibited multiple issues, including use of deprecated ‘pandas’ function, outdated knockout function, and production-envelope plot function, while still scoring well on readability, examples, and clear IO (print and plot features). GPT (37.25) has a similar error on biomass ID and minimal error handling and gene ID validation. DeepSeek suffered from incomplete code, lacked ID verifications, and offered no explanations for its knockout suggestions. Overall, all models would benefit from more rigorous verification of biomass and gene IDs, and parallelization to accelerate multiple knockout evaluations.

Lastly, we evaluated the growth-coupled analysis by prompting LLMs to generate script using COBRApy to determine whether the limonene production through the MVA pathway in iML1515 model is growth-coupled. Among the four LLMs, only Gemini produced fully functional code and achieved the highest score (42.75). It implemented two complementary methods (direct FBA comparison and production-envelope analysis) with plotting and robust error handling for file loading, reaction retrieval and envelope generation, including informative error messages. Claude ranked next (41.75), also providing both direct flux and production-envelope analysis (illustrating feasible combinations of growth and limonene production), and an additional trade-off analysis showing maximum limonene flux across different fixed growth rates; however, it failed at runtime due to incorrect key assumptions about the production envelope’s outputs. DeepSeek and GPT performed moderately (39.75 and 39.5, respectively): DeepSeek generated production-envelope analysis but used an incorrect attribute for the objective function (biomass) and incorrect key assumption for the production envelope’s output, while GPT relied on incorrect biomass ID and applied a medium reset that may induce infeasible growth, leading to erroneous results. In this task, some errors were not fully captured by the evaluation markers.

Overall, across the four flux optimization coding tasks, Gemini produced functional codes for three tasks (Supplementary Table S8) and led most evaluation criteria (Figure 5a), notably error handling, simplicity, correctness, and documentation, achieving the highest consolidated mean score of 42.56 (Figure 5c). Claude placed second (39.75) with two executable solutions; while GPT (36.88) and DeepSeek (36.81) lagged substantially, each succeeding on only one task in this category. At the task level (Figure 5b), correctness exhibited the greatest variability, whereas error handling and efficiency were consistently weak across models. Despite lacking some optimal features (Supplementary Table S8), most LLMs were able to implement the FVSEOF and knockout analysis tasks. However, all models failed the succinate knockout analysis involving the iJR904, largely due to incorrect assumptions for biomass and gene IDs; this suggests that stringent ID verification, especially for models other than iML1515, could improve robustness. Similarly, most models failed the growth-coupled analysis because they assumed an incorrect output key for the production envelope function, indicating a need for explicit key validation aligned to the specific COBRApy function being used. These results highlight that strong average performance does not guarantee output reliability (Figure 5d), indicating a need for scenario-dependent weighted evaluation for different criteria to ensure robustness. Incorporating systematic ID and function key validation is essential for making LLM-assisted workflows consistently reproducible and transferable across GSMs.

### 3.6. Case Study Validation and Overall Performance

To demonstrate the utility of LLMs in a real-world case study, we conducted nine additional validations based on a recent comprehensive evaluation of microbial cell factories capacities using GSM analysis^6^. The first case study involved reading and summarizing the paper^6^. DeepSeek performed best (43.25) (Figure 6a), delivering comprehensive, critical synthesis supported by quantitative examples. GPT followed closely (42.5), with clear structure and strong critique but fewer concrete case studies. Claude (41.75) covered the essentials yet needed sharper limitation analysis and more examples. Gemini performed weakest (40.25), providing a brief, non-elaborative summary that lacked finer details, critique, and case-based evidence. Overall, most LLMs (DeepSeek, GPT, Claude) performed well in paper summarization with well-structured and elaborative key points, except Gemini which lacked elaborative details. All LLMs could benefit from deeper technical specifics, clearer discussion of computational approach limitations or implementation challenges, and richer case-study grounding.

**Figure 6:**
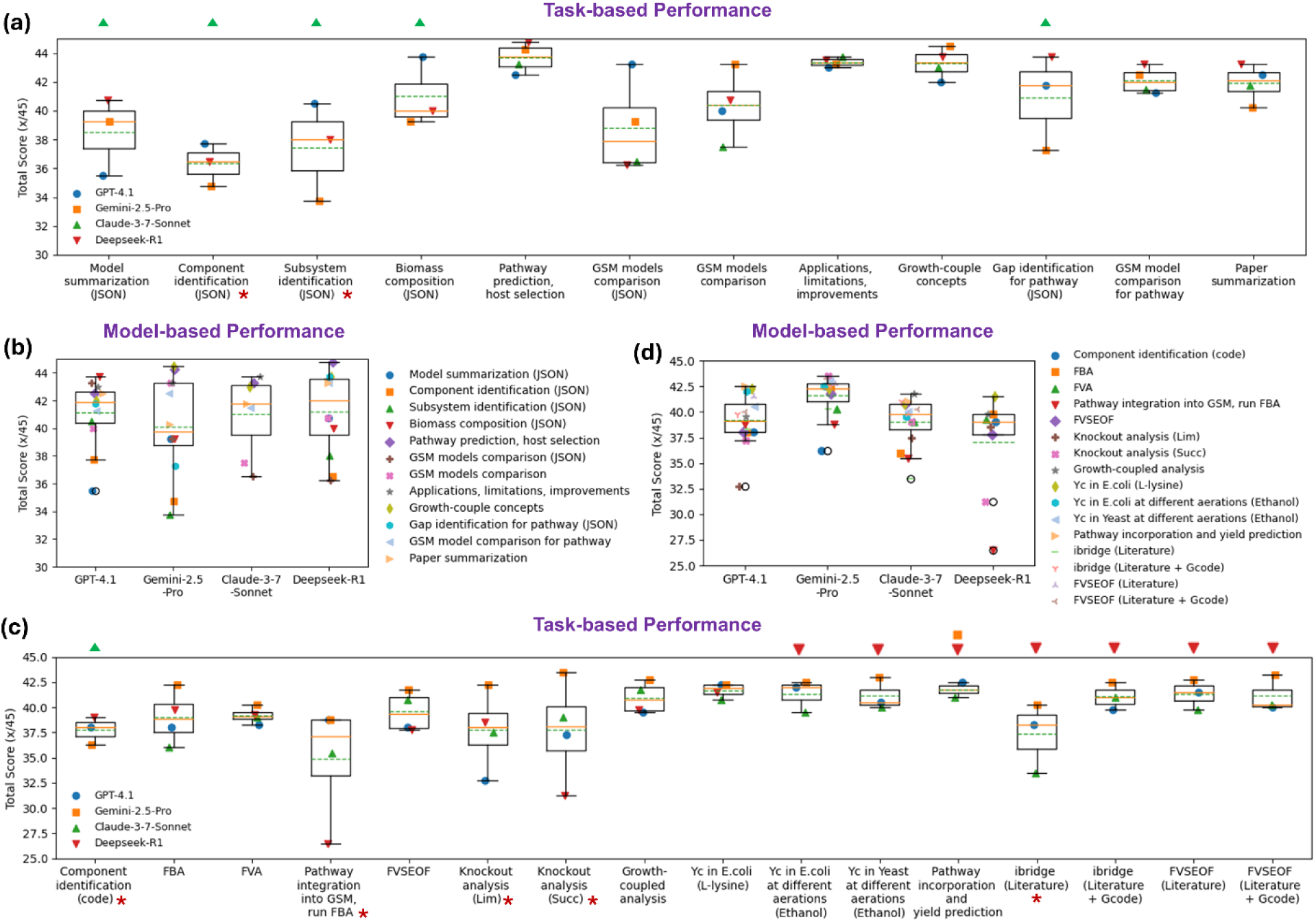
Overall performance across all tasks, including case study validations. (a) Task-based performance for domain tasks (b) Model-based performance for domain tasks (c) Task-based performance for coding/computational tasks (d) Model-based performance for coding/computational tasks.

Next, we selected an L-lysine example from the paper and prompted LLMs to generate code that computes both theoretical yield (YT) and maximum achievable yield (YA) in an *E. coli* GSM using D-glucose under aerobic conditions. All LLMs produced runnable codes, but only Gemini and DeepSeek returned correct yields, while GPT and Claude produced incorrect fluxes due to improper media shutdown (non-carbon sources) and lack of model reset when calculating the different yields. Gemini and GPT achieved the highest score (42.25) (Figure 6c), correctly setting up the constraint for both yields: YT ignores growth and maintenance, and YA accounts for non-growth associated maintenance and enforces biomass viability (10% of maximum growth). In practice, Gemini was more robust; after the initial medium shutdown, Gemini restored key nutrients (Nitrogen and phosphate) and used model context managers, which were omitted by GPT, resulting in GPT’s incorrect final fluxes. DeepSeek followed (41.5), correctly using the context managers for the two yield scenarios and adding a solution optimality check, but lacked safeguards for missing reactions, explicit error messages, and biological commentary. Claude scored 40.75, despite implementing a loopless function consistent with the paper’s methodology^6^, it failed to reset model when switching between yield calculations, leading to incorrect fluxes, and lacked robust checks (missing files, reactions IDs, and solution optimality). LLMs generally demonstrate strong code generation and methodology awareness, but their reliability hinges on robust error handling, correct media configuration, and proper scenario isolation via model reset; lapses in any of these can lead to incorrect results.

Leveraging data provided by the paper, we increased task complexity by prompting the LLMs to compute YT and YA of ethanol in *E. coli* using D-glucose under (i) aerobic (ii) anaerobic and (iii) microaerobic conditions, following the settings described in the paper^6^. Building on earlier insights, we instructed models to retain the medium’s default settings. Nonetheless, Gemini and GPT still disabled all media (exchange bounds); GPT attempted to restore the essential nutrients, yet both models ended up with incorrect values for biomass and YA. Gemini scored highest (42.5) (Figure 6c) for effective use of context managers and solid handling of common issues, though it relied on assumptions for reaction IDs. GPT delivered comparable performance with good error handling of missing reaction IDs but introduced overhead via deep copies and failed to reset model state between yield calculations. Claude, while executable, scored lower (39.5) due to risks such as computing yields by counting carbon atoms, relying on model copies (reduced efficiency), lacking explicit error handling, and assuming an ATPM value, which led to small deviations in YA. DeepSeek failed to generate complete code for this task, as the reasoning required exceeded its available context window, preventing it from handling the task’s complexity. Overall, these results underscore that multi-conditioned computations magnify the importance of strict media adherence, careful model state management, and explicit error handling; small implementation choices can affect both correctness and efficiency.

We next evaluated the same previous computation task using the yeast GSM (yeast850.xml)^58^. In this scenario, all LLMs failed to reproduce the literature reported YT and YA^6^. Gemini achieved the highest score (43) (Figure 6c), demonstrating proper model context management and robust exception handling of missing model files and failed optimizations, heavily praised by the automated ensemble judges, but it incorrectly assumed all IDs without proper verification, resulting in inaccurate yield values. These mistakes were not effectively flagged by the LLM markers, likely reflecting limited yeast-specific knowledge, exposing a critical systemic vulnerability where the ensemble judges share identical training blind spots regarding non-*E. coli* nomenclature. GPT ranked second (40.5), incorporating helper functions for ID checks, yet several key ID issues persisted (e.g., O2 and ethanol), leading to incorrect yields; although it utilized loopless FBA in accordance with the literature, reloaded model in a loop lowered its efficiency score. Claude followed (40), also used loopless FBA, but suffered from inefficient model copying, lacked checks for reaction existence and separate model copy for yield calculations, and disabled certain nutrients, resulting in incorrect outcomes. DeepSeek was not applicable for this case due to context window limitations. Manual verification revealed that, after correcting the errors, all models were able to achieve the same YT, but deviations in YA persisted, potentially attributed to additional constraints applied to the yeast model under microaerobic and anaerobic conditions^77^. This evaluation demonstrated that existing LLMs performed poorly for strains beyond *E. coli* model, and their mistakes were rarely detected by the LLM makers, underscoring the need for human oversight.

Next, we evaluated the capability of pathway integration (specifically the R-mevalonate pathway) into the GSM (iCW773) of *Corynebacterium glutamicum*, a strain known for large-scale amino acid production^78^. The task required LLMs to add the pathway to the model and perform FBA to determine YT, given the GSM JSON file, the three reactions starting from acetyl-CoA, and bioproduction conditions (glucose uptake and O_2_ fluxes). In this case, only GPT and Claude generated functional codes, while Gemini and DeepSeek failed to produce the code files. Both GPT and Claude correctly implemented all steps: adding metabolites and reactions, incorporating exchange reaction, setting constraints, running FBA, and calculating yields. GPT achieved the highest score (42.5) (Figure 6c), with marginal higher efficiency and error handling: efficient use of COBRApy API and checks for existing metabolites to avoid duplicates, though batch addition of metabolites and implementation of try-except blocks for file and key errors could further improve performance. Claude scored 41, with correct implementation and efficient use of cobra functions, but received minor deductions for unused imports and lack of exception handling for file loading or missing reactions. Only GPT and Gemini performed proper markings; the other LLMs failed due to incomplete interpretation of input data, likely caused by context window limitations.

For case studies on metabolic flux optimization, we assessed the LLMs’ abilities to implement the iBridge algorithm based on the literature descriptions^6^, aiming to identify up- and down-regulation targets that enhance production flux (e.g., ethanol in an *E. coli* model using glucose under aerobic conditions). Only Gemini (score: 40.25) (Figure 6c) closely adhered to the iBridge methodology as described in the paper^6^: applying parsimonious FBA (pFBA) for a reference state, generating flux distributions with linear MOMA, calculating covariances, and identifying targets through a metabolite-centric analysis; but failed to generate correct results due to incorrect medium setup. GPT (score: 38.25) implemented only a simplified flux correlation analysis, omitting the core metabolite-centric steps (calculation of sum of covariances and identification of bridge reactions), resulting in a major logical error and divergent results. Claude (score: 33.5) exhibited more pronounced errors: used deprecated DataFrame method, and incorrect normalization and metabolite classification logic. These findings indicate that most LLMs struggle to accurately interpret and implement more complex or domain-specific computational algorithms.

To improve reproducibility, we additionally embedded the relevant GitHub implementation for the iBridge algorithm^79^ aside from the literature and instructed the LLMs to account for the excluded metabolites. Under this setting, Gemini achieved the highest overall score (42.5) (Figure 6c), outperforming others in most evaluation criteria; its implementation was otherwise correct but failed due to COBRApy version-related bug in solver-status checking for biomass feasibility. Claude followed (41), correctly capturing iBridge logic but failing due to a recurring medium reset error. GPT produced outputs like those reported in the literature but scored lower (39.75) due to weaker error handling and insufficient documentation (lacked detailed docstrings). Again, DeepSeek was not applicable for this task. This task demonstrates that providing both literature and relevant code substantially enhances the reproducibility and implementation fidelity of specific algorithms, lowering the barriers for non-experts.

Similar frameworks were applied for evaluating another flux optimization algorithm FVSEOF, using the same ethanol example (glucose, aerobic conditions). With the literature description alone, Gemini (score: 42.75) and GPT (score: 41.5) (Figure 6c) implemented the FVSEOF workflow, utilizing COBRApy functions, proper medium setup, loopless FVA, and Pearson correlation analysis to identify the up- and down-regulation targets, but reproduced only three literature-reported targets. This partial match likely results from failing to reset the model objective to biomass prior to performing FVA under enforced ethanol scanning. Overall, Gemini outperformed GPT in readability, efficiency, error handling, and code documentation. While both implementations were efficient for typical analyses, loopless FVA inside iterative loops can be slow and would benefit from parallelization. Claude (score: 39.75) implemented the core FVSEOF logic and included additional result plotting, but lacks loopless constraints, no efficient use of context managers and objective reset to biomass, resulting in no match to the reported targets. DeepSeek was not applicable in this case and the subsequent case, constrained by its context window limit.

After incorporating both the relevant GitHub implementation^80^ and the literature, Gemini achieved the highest score (43.25) (Figure 6c): produced similar codes with several improvements such as cleaner structure, better handling of biomass constraint and the use of signed flux from Pearson correlation, which is more interpretable; but medium reset could be risky. The model used multiprocessing for FVA like the GitHub code but could leverage solver parallelism. Claude (score: 40.25) also offered a cleaner version of the GitHub code, but initial executions were hampered with errors, primarily due to regression failures by problematic numeric entries; filtering infinite or missing values resolved the issue. Once corrected, both Gemini and Claude reproduced the GitHub results. GPT (score: 40) distilled the complex GitHub workflow while reserving the core concept, but substituted FBA for full FVA for each reaction, and blocked uptakes of other essential nutrients, resulting in only three matched targets as the literature. In this case, we realized a discrepancy between results from the GitHub implementation^80^ and those reported in the literature^6^, which may confuse reproducibility but does not impact the validity of this study. Overall, these findings underscore the importance of consistency and completeness in both literature and code for improved implementation fidelity and demonstrate the potential of LLMs to refactor complex pipelines while preserving core logic.

Upon completing all case studies, we quantified overall performance across all tasks. For domain-specific tasks (Figure 6a), the majority (n=10) achieved an impressive mean score of over 38 out of 45 (∼85% of total score), reflecting strong domain proficiency. The sole exception was component and subsystem identification tasks, which required careful interpretation and targeted retrieval from an extensive GSM JSON file. In the model-based evaluation (Figure 6b), DeepSeek emerged as the leader in domain task performance with a consolidated mean score of 41.21, narrowly ahead of GPT (41.15), Claude (41.04), and Gemini (40.13). Notably, Claude failed five tasks involving GSM JSON input, primarily due to context window constraints. For coding-focused tasks (Figure 6c), five out of 16 tasks achieved mean scores below 38, including component identification, pathway integration, two knockout analyses, and iBridge (literature only). Gemini ranked first in coding tasks (41.62) (Figure 6d), followed by GPT (39.20), Claude (39.02), and DeepSeek (37.03). Execution failures were most frequently observed for coding tasks, particularly with DeepSeek (7 failures), Claude (1), and Gemini (1), primarily attributed to context window limitations. Collectively, these findings underscore the models’ overall robustness in domain scientific reasoning and coding, while highlighting the decisive bottleneck of context limit for handling complex tasks and large data inputs, especially GSM. The results also demonstrate that, with careful configuration, LLMs can achieve high accuracy and reliability across diverse sets of scientific workflows.

## 4. Discussions

In this study, we systematically benchmarked four leading LLMs (GPT-4.1, Gemini-2.5-Pro, Claude-3-7-Sonnet, and DeepSeek-R1), assessing their capabilities across four core areas central to the application of GSMs in metabolic engineering: domain knowledge, metabolic flux prediction, metabolic pathway construction, and metabolic flux optimization. Our evaluation encompassed 28 distinct tasks through a total of 112 *in silico* experimental runs, revealing distinct strengths and limitations. Notably, DeepSeek and GPT demonstrated superior performance in domain knowledge tasks, benefiting from their robust factual grounding and reasoning process. Gemini outperformed others in coding-centric tasks, producing more readable, better-documented code with stronger error handling, despite occasional biological-context errors. GPT’s ability to handle tasks requiring a high context window limit proved valuable, as evidenced by its resilient against context window failures that affected Claude and DeepSeek. However, all models struggled with more complex tasks such as pathway model construction and metabolic flux optimization, indicating that human oversight remains essential. Additionally, issues such as metabolite and biomass ID mismatches, handling strains beyond *E. coli*, and limitations with large GSM JSON files highlighted practical barriers that must be addressed for broader utility. Rather than viewing context window disruptions strictly as an intrinsic hardware limitation of modern architectures, these failure modes uncover a deeper structural flaw in conversational prompt engineering. Forcing a language model to perform quasi-string matching over a 150,000-line structural network is fundamentally unscalable. To transition LLMs from fragile conversational “copilots” to automated, production-ready metabolic engineering pipelines, the design paradigm must shift entirely away from monolithic text ingestion. Future workflows must deploy token-lean agentic architectures; in this decoupled design, the LLM does not ingest the raw model, but instead acts as an orchestrator that interacts with the network programmatically via an abstracted tool-use layer—writing discrete python code snippets to inspect localized matrix rows (model.reactions. get_by_id()) and processing data iteratively in a sandboxed runtime.

To enhance auditability of the benchmarking workflow, we proposed and demonstrated the utilities and advantages of applying multi-LLM auto-evaluation approach. Operationally, this approach is highly scalable; models can process and score large batches of responses in parallel. It is both cost- and time-efficient; automated evaluation is substantially less resource-intensive than expert manual grading especially given the complexity of multiple criteria and scales. Additionally, this method is highly reproducible; consistent application of rubric and inputs minimizes human bias and inter-grader variability. By leveraging diverse models, we reduce single-model bias; consensus and consolidated mean scores improve accuracy and stability whereas disagreement flags help identify ambiguous cases for further human review. As different models excel at distinct tasks (e.g., scientific explanation or code inspection), combining them yields more balanced judgements. Additionally, variance across models serves as an indicator of inter-rater reliability for each criterion. While all criteria were weighed equally in this study, applying scenario-specific weighting could further strengthen the evaluation; enhancing the reliability and interpretability of the outputs and improving the robustness of conclusions. Taken together, the proposed multi-LLM auto-evaluation framework delivers a robust, transparent, and reliable solution, that is highly applicable and easily adaptable for large-scale benchmarking across diverse domains, from education and research to industry.

To assess LLM error-detection capability on complex GSM files, we introduced two subtle, yet consequential faults representative of issues encountered in practice: (i) a stoichiometric sign error that produces mass/charge imbalance, and (ii) a missing reaction. We assessed multiple analysis strategies, including prompt specificity (general vs targeted prompts), text-based inspection vs code-driven analysis, and cross-checking against a reference model file (iML1515.json). In the sign-error scenario (reaction id: PGI; name: Glucose-6-phosphate isomerase), the correct stoichiometry for the PGI reaction is g6p_c_ : - 1, f6p_c_ : + 1; whereas the faulty version is g6p_c_ : - 1, f6p_c_ : - 1. Under general prompting, approaches relying on text interpretation, code inspection, or reference file comparison (text or code-based) failed to recover the true fault. Robust detection was achieved only when the workflow explicitly invoked COBRApy function to identify mass balance issues, with boundaries reactions and the biomass reaction excluded from the imbalance screen. For the missing-reaction scenario (omitted reaction id: TPI; name: triose phosphate isomerase; expected equation dhap_c_ ↔ g3p_c_), “blind search” general prompting, using either text-based interpretation or code-driven analysis, flagged several dead-end metabolites but failed to identify the correct omitted reaction. In contrast, a targeted prompt with guided steps to perform pathway-completeness analysis, and code-based reference model comparison successfully localized the intended omission. Collectively, these error detection analyses uncover the limitations of LLMs in reliably handling large GSM content and demonstrate that general prompting is insufficient for reliable error localization in large GSMs, whereas prompts reframed with domain-informed constraints and tool-assisted procedures can substantially improve detection performance (Refer Supplementary Table S14 for the list of prompts used for analyses)

This study systematically unraveled recurrent failures modes and both task- and model-specific limitations when deploying LLMs within GSM workflows, providing a robust foundation for articulating best practices in the application. Meanwhile, we highlight the significant potential of LLMs to advance metabolic engineering: domain knowledge, pathway suggestions, and metabolic flux prediction; models such as DeepSeek and GPT reliably deliver domain knowledge, while Gemini’s coding capabilities lower technical barriers for non-experts. Despite the challenges encountered such as context window limits, incorrect ID assumptions, and limited support for strains beyond E. coli, LLMs are well positioned as “copilots” for GSM-based analyses, enhancing both productivity and understanding. For novices, LLMs support the steep GSM learning curve by providing accessible domain knowledge and enabling simple coding-driven analyses. For experts, it can accelerate routine analyses when augmented with effective prompt engineering and verification. The insights from this benchmarking study inform the next phase: developing more reliable, accessible, and automation-ready agentic workflows for pathway and strain design, ultimately further lowering barriers to entry and accelerating innovation in metabolic engineering.

## 5. Conclusions

Overall, this comprehensive benchmarking study underscores that no single LLM is universally optimal across all facets of metabolic modelling. DeepSeek and GPT are the preferred choices for domain knowledge retrieval, while Gemini excels in coding and error detection, albeit with some contextual limitations. The frequent failures due to context window limits, as well as difficulties in complex pathway construction and optimization tasks, signal crucial areas for improvement in future AI development in the context of GSMs. Integrating multi-model orchestration and advancing context window capabilities could be most beneficial to more reliably pair the precision of AI with human expertise to streamline and expedite metabolic engineering workflows.

## CRediT author contribution statement

Y.J.W.: Conceptualization, Methodology, Formal analysis, Validation, Writing-Original Draft, Writing-Review & Editing. C.P.K.P.: Conceptualization, Writing-Review & Editing. W.L.S.: Conceptualization, Supervision, Writing-Review & Editing. P.C.L.: Conceptualization, Resources, Writing-Review & Editing, Supervision, Project administration, Funding acquisition.

## Acknowledgments

The authors gratefully acknowledge the funding support by the National Centre for Engineering Biology (NRF-MSG-2023-0003) by The National Research Foundation, Singapore. The authors thank Liao Wenjun, Ji Yutong, Yossa Dwi Hartono, and Lim Sook Wei for their contributions to ideation and discussion.

