## Supplementary Materials for "Comprehensive evaluation of LLM capabilities for interpretation and analysis of genome-scale metabolic models in metabolic engineering"

*Corresponding author

**Table S1:** rubric-based evaluation metrics used in multi-model auto-evaluation for domain tasks

| **Metric/score** | **1 (Poor)** | **2 (Fair)** | **3 (Adequate)** | **4 (Good)** | **5 (Excellent/Outstanding)** |
|---|---|---|---|---|---|
| **Accuracy** | Major errors, mostly incorrect | Several errors, some correct info | Minor errors, mostly correct | Accurate with only trivial errors | All information is fully correct and precise |
| **Completeness** | Major gaps, large portions missing | Some key points missing | Covers most key aspects but some minor omissions | Covers all essential points, minor details missing | Fully complete, all aspects thoroughly addressed |
| **Relevance** | Largely off-topic, irrelevant | Some relevant points but much off-topic | Mostly relevant, some off-topic information | Nearly all content is relevant | Entirely focused and directly relevant to GSM |
| **Depth/Details** | Superficial, lacks detail | Limited depth, only basic points | Some technical details, not thorough | Good depth, clear explanations and supporting details | In-depth, substantial technical detail and insightful explanations |
| **Clarity** | Unclear, confusing | Frequently unclear, hard to follow | Generally understandable, some unclear sections | Well explained, minor lapses in clarity | Exceptionally clear, easy to follow, no ambiguity |
| **Organization** | Disorganized, illogical | Some structure but poorly organized | Structure is present but could be improved | Well organized, logical flow | Exceptionally well structured and logically organized |
| **Conciseness** | Overly verbose, | Some redundancy, | Adequately concise, some | Mostly succinct, | Highly succinct, clear, no |

| | | | | | |
|---|---|---|---|---|---|
| | much repetition | occasionally wordy | minor redundancy | minimal extraneous info | unnecessary information |
| **Critical Analysis** | No analysis, only repeats facts | Minimal critical thought, weak analysis | Some analysis, but limited depth | Sound critical evaluation, identifies strengths/weaknesses | Deep, insightful analysis, clearly evaluates limitations and alternatives |
| **Examples/Analogies** | None provided | Irrelevant or unclear examples | Somewhat relevant examples, not fully developed | Good, relevant examples or analogies, may lack depth | Highly effective and illuminating examples or analogies, well integrated with explanation |

**Table S2:** rubric-based evaluation metrics used in multi-model auto-evaluation for computational/coding tasks

| Metric/score | 1 (Poor) | 2 (Fair) | 3 (Adequate) | 4 (Good) | 5 (Excellent/Outstanding) |
|---|---|---|---|---|---|
| **Readability/Style** | Poor formatting, hard to read | Inconsistent style, partially readable | Moderately readable, can be improved | Clear and consistent style, easy to follow | Exceptionally readable and well formatted |
| **Correctness (code)** | Code incorrect/does not run | Significant bugs or logical errors, results mostly wrong | Minor errors, results are mostly correct | Results correct with only trivial issues | Fully correct execution and results, robust code |
| **Efficiency** | Highly inefficient, unnecessary complexity | Some inefficiency | Acceptable efficiency | Efficient, well optimized | Highly optimized, best practice efficiency |
| **Simplicity** | Overly complex, hard to understand | Somewhat complex, not streamlined | Adequately simple, but could be clearer | Simple, clear solutions | Elegant, minimal, and highly understandable solutions |
| **Error Handling** | No error handling | Minimal checks, fragile code | Basic error handling, covers common cases | Robust against most errors | Anticipates and manages all likely errors gracefully |
| **Example Provided** | No example | Example incorrect or unrelated | Example included but unclear or incomplete | Good, relevant example, mostly complete | Excellent, clear, relevant, and comprehensive example |
| **Clear IO** | Unclear, missing input/output | Partially clear, some confusion | Input/output mostly clear, some ambiguity | Clear input/output, minor possible improvements | Extremely clear and well documented input/output |

| | | | | | |
|---|---|---|---|---|---|
| **Documentation/Comments** | None present | Minimal or unclear comments | Some documentation, could be more informative | Well-documented, most choices explained | Exceptionally well-documented, every step and rationale is clear |
| **Biological Mapping** | No biological connection, not GSM-relevant | Minimal biological context, unclear mapping | Some mapping to biological scenario, partially relevant | Good mapping, interprets computational work biologically | Direct, insightful mapping to relevant biological questions or scenarios |

```
Prompt = f"""
You are a marker. Use the following rubric to grade the answer based on the question.

Rubric:
{rubric_text}

Q & A:
{LLM_answer}

Return filename = {result_file}, and a table with the score for each criterion and a short feedback/rationale.
"""
```

**Figure S1:** Grading prompt template used for rubric-based marking. The prompt instructs the LLM to act as a marker (domain tasks), apply the provided rubric to evaluate the Q & A (query and answer) for different tasks, and output the result file plus a per-criterion scoring table supported with brief feedback/rationale. For coding tasks, we adapted the marker into code marker to reflect its more specific role to better capture code implementation details.

**Table S3:** Performance evaluation (task-based and model-based summary table) for domain knowledge tasks

| **Evaluation tasks** | **GPT** | **Gemini** | **Claude** | **DeepSeek** |
|---|---|---|---|---|
| Model overview + JSON structure tree | Code-based extraction; functional code to provide concise summary and a visual overview of model's structure but limited biological insights/critical analysis **(Score: 35.5)** | Covered major sections + counts + clear JSON hierarchy tree with examples, purely descriptive, no evaluation of model limitations and biological context. **(Score: 39.25)** | **Not applicable** (context window limitation) | Code-based extraction; functional to provide concise summary and visual structure with examples, but fewer biological insights/critical analysis **(Score: 40.75)** |
| Component identification (compound → | Incomplete/incorrect reaction lists; pathway errors propagate | Missing list of major reactions, incomplete/incorrect reaction list, lack | **Not applicable** (context window limitation) | Incomplete (minor)/incorrect reaction list; purely factual report; lack |

| | | | | |
|---|---|---|---|---|
| reactions + pathway) | **(Score: 37.75)** | details explanation of pathways, limited metabolic significance evaluation. **(Score: 34.75)** | | metabolic significance evaluation **(Score: 36.5)** |
| Code-based component identification (accuracy-focused) | Best functional code + accurate results + step-by-step explanation; minor deductions for limited error handling **(Score: 38)** | Weakest with truncated JSON input strings and limited examples **(Score: 36.25)** | **Not applicable** (context window limitation) | Errors: incomplete file loading; unvalidated pathway identifier assumption; still efficient with clear IO + example provided + some biological mapping **(Score: 39)** |
| Subsystem reaction/pathway summarization | Good breadth/depth, but omissions + inaccuracies/inference without evidence **(Score: 40.5)** | Only factual listing, lacks reaction function, pathway context, metabolic-role analysis **(Score: 33.75)** | **Not applicable** (context window limitation) | Good breadth/depth, but omissions + inaccuracies/inference without evidence **(Score: 38)** |
| Biomass objective explanation (role, composition, WT vs core) | Best overall: thorough explanation + implementation details + stoichiometry examples + comparison table; could add limitations (static/condition-dependent) **(Score: 43.75)** | Generally accurate explanation; less thorough than GPT; less critical evaluation **(Score: 39.25)** | **Not applicable** (context window limitation) | Generally accurate explanation; less thorough than GPT; less critical evaluation **(Score: 40)** |
| Biosynthetic pathway prediction, host selection | Accurate MVA genes/enzymes; good bottlenecks + solutions **(Score: 42.5)** | Accurate pathway + host suggestions + bottleneck analysis + modifications **(Score: 44.25)** | Accurate MVA genes/enzymes; good bottlenecks + solutions **(Score: 43.25)** | Near-prefect: accurate + concrete strain designs (E. coli + yeast) **(Score: 44.75)** |
| Compare two GSMs with parsed inputs (E. coli vs yeast) | Reads both; covers major and minor differences; structured table + conclusions **(Score: 43.25)** | Reads both but limited scope; mainly major pathways; lacks minor pathways + deeper critique/analogies **(Score: 39.25)** | Limited: only interpreted one GSM → fails true comparison **(Score: 36.5)** | Limited: only interpreted one GSM → fails true comparison **(Score: 36.25)** |
| Compare two GSMs without providing GSM inputs (knowledge-based comparison) | Minor but significant quantitative deviations (esp. yeast8) + limited implications/critical analysis **(Score: 40)** | Best overall: good implications + engineering applications in conclusion **(Score: 43.25)** | Lowest: weaker completeness/depth/critical analysis/examples **(Score: 37.5)** | Minor but significant quantitative deviations (esp. yeast8) + limited implications/critical analysis **(Score: 40.75)** |
| Role of GSM: applications, | High performance: in-depth information | High performance: in-depth information | High performance: in-depth | High performance: in-depth information |

| | | | | |
|---|---|---|---|---|
| limitations, and improvements | supported with examples and references for applications + limitations + improvements **(Score: 43)** | supported with examples and references for applications + limitations + improvements **(Score: 43.25)** | information supported with examples and references for applications + limitations + improvements **(Score: 43.75)** | supported with examples and references for applications + limitations + improvements **(Score: 43.5)** |
| Growth-coupling bioproduction concepts | Expert-level; could add more algorithm/tool trade-offs + mechanistic details **(Score: 42)** | Best overall; highly effective examples integrated seamlessly into explanations **(Score: 44.5)** | Expert-level; could add more algorithm/tool trade-offs + mechanistic details **(Score: 43)** | Expert-level; could add more algorithm/tool trade-offs + mechanistic details **(Score: 43.75)** |
| **Summary** | Top performers for four tasks; strong structured reasoning for core GSM concepts; perform best on GSM-to-GSM comparison when partial model fields are provided; In GSM-input reasoning tasks, tends to show omissions or inference without direct evidence; accuracy improves when switching to code-grounded extraction but still benefit from stronger explicit validation/critical analysis. **(Mean: 40.625)** | Top performers for two tasks; high variations among tasks; response more factual listing, lack critical evaluations and under-deliver on deeper functional interpretation in some cases. **(Mean: 39.775)** | Top performer for one task; high-quality domain explanations when not input-limited; Primary limitation is context window for GSM-embedded prompts; making several GSM-input reasoning tasks not feasible; In knowledge-only comparisons, tend to be weaker on depth/critical analysis/examples/ analogies relative to others. **(Mean: 40.8)** | Top performers for three tasks; very strong on pathway design/domain prompts; In GSM-input reasoning tasks, more prone to incompleteness and unvalidated assumptions, plus omissions/incorrect inclusions when identifying reactions/pathways from embedded model content; often relies on a code-first approach but may implement it imperfectly. **(Mean: 40.325)** |
| *Across the domain-knowledge tasks, LLMs consistently showed strong fundamental understanding of GSM concepts (e.g., biomass objectives, growth-coupling, pathway design) and could often provide clear, well-structured explanations. The main limitation was reliability on GSM-input reasoning: models commonly omitted key reactions/pathways, made unverified assumptions about identifiers, and sometimes inferred biological interpretations without direct evidence from the provided data. Careful validation (via code-based extraction and human review) remains essential. Constrained by the context window limitation, comparing multiple GSM files might be infeasible. | | | | |

**Table S4:** Performance evaluation of LLMs across metabolic flux prediction tasks.

| Evaluation tasks | GPT | Gemini | Claude | DeepSeek |
|---|---|---|---|---|
| FBA on GSM (e.g., iML1515.json): run FBA + identify top 10 | High correctness: proper COBRApy model loading, FBA execution, prints objective (biomass), | High correctness + very clear explanations and formatting. Best error handling: robust | High correctness: appropriate COBRApy use; clear, well-formatted | High correctness and readability; minimal error handling (e.g., solution optimality |

| | | | | |
|---|---|---|---|---|
| highest-flux reactions | sorts/prints top 10 reactions with details. Strong readability/simplicity. Weaker error handling (omits robust safeguards), making code more fragile. **(Score: 38)** | try/except for missing file, solution status, and general exceptions with helpful messages/instructions. Produces correct results. **(Score: 42.25)** | explanations; elegant/minimal solution. Weaker error handling (generally omits safeguards). Produces correct results. **(Score: 36)** | check). Produces correct results. **(Score: 39.75)** |
| FVA (identify top 10 redundant reactions with the largest flux range | Use COBRApy FVA function, simple and highly readable, but no error handling. **(Score: 38.25)** | Exceptional documentation, print details from model, basic error handling (e.g., file loading) **(Score: 40.25)** | Solution is efficient and good use of pandas for data manipulation, but no error handling. **(Score: 39)** | Code/solution is straightforward, minimal, and highly understandable, no error handling **(Score: 39.25)** |
| **Summary** | Reliable on correctness/simplicity/efficiency; tends to lag mainly on error handling. **(Mean: 38.125)** | Top overall in both tasks, especially in error handling and documentation/comments. **(Mean: 41.25)** | Reliable correctness/readability; lower means mainly due to limited error handling. **(Mean: 37.5)** | Tied 2nd across tasks; strong correctness/readability but minimal error handling. **(Mean: 39.5)** |
| *LLMs deliver well-structured, correct implementation of FBA and FVA analyses using COBRApy and pandas for data manipulation, with exceptionally readable and clear comments. The main weakness is limited error handling, could be improved by adding more robust error handling for file operations and optimization failures. | | | | |

**Table S5:** Implementation evaluation of LLMs across metabolic flux prediction tasks and the identified optimal features.

| Task | GPT | Gemini | Claude | DeepSeek | Optimal Features |
|---|---|---|---|---|---|
| FBA on GSM (identify top 10 reactions with the highest flux) | ● | ● | ● | ● | ▪ 'cobra' package installed<br>▪ Built-in FBA function from cobrapy package<br>▪ Try/except handling for missing file, solution status.<br>▪ Tabular report |
| FVA (identify top 10 redundant reactions with the largest flux range | ● | ● | ● | ● | ▪ 'cobra' and 'pandas' packages installed<br>▪ Built-in FVA function from cobrapy package<br>▪ Solution optimality checks<br>▪ Tabular report |
| *The solid circle indicates that the model generated functional code that executed without errors, whereas the unfilled circle denotes that the generated codes encountered at least one error during execution. | | | | | |

**Table S6:** Key insights for task-based and model-based summary table for metabolic pathway construction tasks

| Evaluation tasks | GPT | Gemini | Claude | DeepSeek |
|---|---|---|---|---|
| Gap identification in GSM for pathway (limonene): read JSON; propose gap-filling reactions/enzymes | Strong: covers heterologous additions to native MEP, also considers MVA alternative; good analysis of precursor limits/competition **(Score: 41.75)** | Weaker: focuses only on MEP; identifies missing gap + required enzymes but minimal reaction/EC details and limited evaluation/alternatives **(Score: 37.25)** | **Not applicable with GSM JSON input** (context window limitation) | Best: most comprehensive (gaps, candidate reactions, heterologous enzymes, considerations, sample pathway addition), but only MEP (misses MVA alternative) **(Score: 43.75)** |
| Strain comparison/selection (knowledge-based; no GSM files) | Fully addresses gaps, comparison, and improvement strategies; lacks technical methods (FBA, OptKnock); lacks specific analogies **(Score: 41.25)** | Includes industrial vs theoretical considerations; could have included more quantitative details and data; lacks specific case studies **(Score: 42.5)** | Could include more quantitative or pathway-specific details; specific genes for knockout are omitted **(Score: 41.5)** | Best: deepest technical analysis (compartmentalization/cofactors), mentions FBA/FVA, uses quantitative literature examples + specific knockouts **(Score: 43.25)** |
| Pathway Integration + FBA (propose MVA route + write code to add to GSM model and optimize pathway (limonene) | Only model with fully functional code (adds full pathway + runs flux analysis); missing robust validation checks lowered score **(Score: 38.75)** | Proposes correct pathway, comparable score due to better readability/error-handling/commence, but code had implementation errors (e.g., assume wrong metabolite IDs, incomplete reactions) **(Score: 38.75)** | Proposes correct pathway but code runs with errors (wrong metabolite IDs, stoichiometric balancing) **(Score: 35.5)** | Weakest: Proposes correct pathway but generates incomplete code, many syntax/definition errors, little error handlings **(Score: 26.5)** |
| **Summary** | Domain tasks are strong; and strong for the pathway integration coding tasks; best overall when combining domain + coding across model-construction tasks). Key limitation: insufficient validation/error handling for IDs in coding integration. **(Mean: 40.583)** | Large performance variance across tasks; domain tasks comparatively weaker, yet coding score comparable to GPT mainly due to better documentation/error handling despite correctness issues. **(Mean: 39.5)** | Strong on domain-focused tasks but limited for GSM ingestion and extended multi-step workflows; weaker for pathway integration coding task. **(Mean: 38.5)** | Best on domain-focused tasks, but weakest on coding integration; suboptimal for long-context GSM ingestion and complex multi-step coding. **(Mean: 37.833)** |
| *Across these pathway-construction tasks, LLMs performed strongly on domain reasoning: identifying pathway gaps, comparing hosts, and proposing biologically sensible yield-improvements strategies; DeepSeek generally provides the most comprehensive technical analyses and examples. The main limitation was reliability in GSM | | | | |

| |
|---|
| pathway integration code: most LLMs produced incomplete or incorrect implementations (ID/stoichiometry/syntax issues and insufficient validation), only GPT generated fully executable code for adding the pathway and running FBA. |

**Table S7:** Performance evaluation (task-based and model-based summary table) for metabolic flux optimization tasks.

| Evaluation tasks | GPT | Gemini | Claude | DeepSeek |
|---|---|---|---|---|
| FVSEOF for succinate in E. coli iML1515 (standardized method: enforced levels → FVA min/max → correlate enforced level vs min flux → top ±10 targets | **(Score: 38.0)** Fast/concise; used slim_optimize() to compute max target flux efficiently. Weaknesses: no biomass viability consideration, used full model copies (not context managers), sparse comments. | **(Score: 41.75)** Correct use of context managers for temporary constraints, loopless FVA, enforced biomass viability, handled NaNs in correlation, strong comments. Downside: loopless FVA slowed execution. | **(Score: 40.75)** Used context managers; enforced biomass viability; improved efficiency by parallel FVA via processes (4 cores). Added plots (correlation distribution, flux profiles) + saved CSV outputs. | **(Score: 37.75)** Key flaw: did not isolate constraints across iterations (no copy/context manager) → distorted results. Omitted infeasibilities check after FVA; produced NaNs for downregulation targets. |
| Single-knockout analysis to improve product yield (e.g., limonene) while maintaining 10% max biomass | **(Score: 32.75)** Non-functional code due to misuse of cobra.flux_analysis.single_gene_deletion (wrong key assumptions) + redundant model loads. Had basic biomass/product checks but no solver optimality/infeasibility check. | **(Score: 42.25)** Production-quality: excellent docs, informative errors/prints, parallelized knockout runs, safe constraint handling via context managers. Minor improvement: verify reaction IDs before use. | **(Score: 37.5)** Validated IDs + solution optimality; did single gene and reaction deletions via manual loop. Extra: CSV outputs and plots. | **(Score: 38.5)** Included solution optimality check; but inefficient manual loop over genes; omitted wild-type comparison for improvement validation; lacked reaction ID verification. |
| Succinate knockout workflow in E. coli iJR904 (anaerobic; suggest KOs + run FBA for biomass-opt + succinate-opt at 10% biomass + implement suggested KOs) | **(Score: 37.25)** Followed intended workflow but failed at runtime due to incorrect biomass reaction ID; minimal error handling and weak gene ID validation. | **(Score: 43.5)** Strong overall (safe context managers; robust model loading and solution-status checks) but still assumed wrong biomass ID and lacked gene ID validation → execution failure. | **(Score: 39.0)** Multiple issues: deprecated pandas' function, outdated knockout function, production-envelope plotting function; still strong on readability/examples and clear I/O (prints/plots). Also failed due to incorrect biomass ID. | Failed: incomplete code; lacked ID verifications; provided no explanations for knockout suggestions. Also failed due to biomass ID issue. |
| Growth-coupled analysis: limonene via MVA pathway in iML1515 (is | **(Score: 39.5)** Used incorrect biomass ID; medium reset may cause infeasible growth → | **(Score: 42.75)** The only fully functional solution. Implemented two methods (direct FBA | **(Score: 41.75)** Provided direct flux + production-envelope analysis plus extra trade-off | **(Score: 39.75)** Implemented production-envelope but used incorrect biomass objective |

| | | | | |
|---|---|---|---|---|
| production growth-coupled?) | erroneous results. Some errors not fully caught by markers. | comparison + production-envelope) with plotting and robust error handling and file loading, reaction retrieval, envelope generation (informative error messages). | analysis (max limonene across fixed growth rates). Runtime failure due to incorrect key assumptions about production-envelope outputs. | attribute and wrong key assumption for envelope outputs → failure. |
| **Summary** | **(Mean: 36.88)** Succeeded on 1/4 tasks in this category; errors often tied to incorrect biomass IDs and weak validation/handling | **(Mean: 42.56)** Functional code in 3/4 tasks; strongest in error handling, simplicity, correctness, documentation. | **(Mean: 39.75)** 2/4 executable solutions; generally readable with helpful outputs; runtime issues frequently from incorrect function/output-key assumptions and outdated calls. | **(Mean: 36.81)** Succeeded on 1/4 tasks; common gaps: constraint isolation, ID verification, completeness/explanations. |
| *Across models (reliability notes/improvements): add explicit try/except for file I/O, reaction IDs, solver errors; check solver optimality after FBA/FVA; validate NaN correlations; rigorously verify biomass/gene IDs (especially beyond iML1515); validate COBRApy function output keys (notably production-envelope); parallelize multi-knockout runs where possible. | | | | |

**Table S8:** Implementation evaluation (task-based and model-based summary table) for metabolic flux optimization tasks and the identified optimal features.

| Task | GPT | Gemini | Claude | DeepSeek | Optimal Features |
|---|---|---|---|---|---|
| FVSEOF | ● | ● | ● | ○ | ▪ slim_optimize()<br>▪ Parallelized FVA<br>▪ Context managers<br>▪ NaN handling<br>▪ Biomass viability |
| Knockout analysis (Lim) | ○ | ● | ● | ● | ▪ Built-in function vs manual loop<br>▪ Parallelized knockout runs<br>▪ Output key assumption for the built-in function<br>▪ Reaction IDs assumption and verification<br>▪ Solution optimality checks |
| Knockout analysis (Succ) | ○ | ○ | ○ | ○ | ▪ Biomass and genes IDs verifications<br>▪ Parallel knockout for speed |

| | | | | | |
|---|---|---|---|---|---|
| | | | | | ▪ Version-dependent built-in functions (knockout and plot)<br>▪ Tabular reports |
| Growth-coupled analysis | ○ | ● | ○ | ○ | ▪ Include direct flux, production envelope, and trade-off analyses<br>▪ Output key verification for production-envelope function<br>▪ Biomass ID verification<br>▪ Careful medium reset |
| *The solid circle indicates that the model generated functional code that executed without errors, whereas the unfilled circle denotes that the generated codes encountered at least one error during execution. | | | | | |

**Table S9:** Performance evaluation (task-based and model-based summary table) for case studies derived from Kim et al. (2025).

| Evaluation tasks | GPT | Gemini | Claude | DeepSeek |
|---|---|---|---|---|
| read and summarize literature | Elaborative, clear sections; good critique of limitations and future directions. Minor improvement: add specific case studies/examples **(Score: 42.5)** | Lowest: brief, non-elaborative; lacks finer details, critical analysis of limitations, and specific case studies. **(Score: 40.25)** | Covers major aspects; needs stronger critical evaluation of limitations and more concrete examples. **(Score: 41.75)** | Best: thorough on objectives, methods, findings, implications; strong strengths/limitations critique; supported with quantitative case studies. **(Score: 43.25)** |
| Compute YT & YA for L-lysine in E. coli GSM; D-glucose, aerobic | Runnable but incorrect yield values due to minor media configuration issue (turning off media) and lack of model reset between yield calculations **(Score: 42.25)** | Runnable and correct yield values; correct constraint setup for YT (ignored growth/maintenance) and YA (including NGAM; biomass variability at 10% max growth); used model context managers. **(Score: 42.25)** | Runnable but incorrect flux due to missing model reset between scenarios; implemented loopless function consistent with paper; limited robustness (missing files/IDs handling; solution optimality checks) **(Score: 40.75)** | Runnable and correct yield values; used context managers across scenarios; added solution optimality check; weaker on safeguards for missing reactions, explicit error messages, and biological-context comments. **(Score: 41.5)** |
| Compute YT & YA for ethanol under aerobic/anaerobic/ microaerobic conditions (use default medium setting) | **(Score: 42)** Comparable performance with good error handling for missing reaction IDs, but incorrect biomass and YA; disabled media and attempted to restore essentials; used deep | **(Score: 42.5)** Highest; effective context managers and handling of common issues; still disabled media despite instruction; relied on assumed reaction IDs; ended with | **(Score: 39.5)** Executable but risk-prone: computed yields by carbon counting; relied on model copies (less efficient); lacked explicit error handling; assumed | Failed to generate complete code due to context-window limitation; could not handle added complexity |

| | | | | |
|---|---|---|---|---|
| | copies (overhead) and failed to reset model state between yield calculations. | incorrect biomass and YA | ATPM, causing small YA deviations. | |
| Yeast GSM computation to reproduce YT & YA for ethanol | **(Score: 40.5)** Used helper functions for ID checks, but key ID issues persisted (e.g., O2, ethanol), causing incorrect yields. Used loopless FBA (as literature) but inefficiently reloaded the model in the loop (lower efficiency) | **(Score: 43)** Good context management; handled missing model files & failed optimizations. However assumed IDs without verification, causing inaccurate yields. *Errors not caught by markers (probably limited yeast-strain knowledge) | **(Score: 40)** Used loopless FBA, but inefficient model copying; missing checks for reaction existence/separate model copies for different yield calculations; disabled certain nutrients, causing incorrect outputs. | Not appliable (context-window limitation) |
| Pathway integration into C. glutamicum GSM: add R-mevalonate + FBA for YT (given GSM JSON file, three reactions for the pathway, and bioproduction conditions) | (Score: 42.5) Generated functional code; correctly added metabolites/reactions + exchange, set constraints, ran FBA, computed yields. Good efficiency and error handling (good Cobrapy API usage; checks to avoid duplicate metabolites); could improve with batch metabolite addition + try/except for file/key errors | Failed to produce code files | (Score: 41) Generated functional code; correctly implemented full workflow; efficient cobra usage. Minor deductions: unused imports and lack of exception handling for file loading/missing reactions. | Failed to produce code files |
| Implement iBridge algorithm based on literature | **(Score: 38.25)** Implemented simplified flux-correlation approach; omitted core iBridge metabolite-centric steps (sum of covariances, bridge reactions) → major logical error and divergent results. | **(Score: 40.25)** Closest to literature: reference pFBA, generate states via linear MOMA, compute covariances, metabolite-centric target identification; but incorrect medium setup → wrong results | **(Score: 33.5)** More pronounced issues: deprecated DataFrame method; incorrect normalization and metabolite classification logic. | Not applicable (context window limitation) |
| Implement iBridge with literature + GitHub code embedded | **(Score: 39.75)** Produced literature-like outputs but lost points for weaker error handling and insufficient documentation/comments (missing full docstrings) | **(Score: 42.5)** Implementation mostly correct; major bug in checking solver status for biomass feasibility (Cobrapy versioning issue), otherwise would have produced correct results. | **(Score: 41)** Correct iBridge logic, but failed due to recurring error during medium reset; otherwise results accurate | Not applicable |

| | | | | |
|---|---|---|---|---|
| FVSEOF from literature only | **(Score: 41.5)** Implemented FVSEOF workflow (proper medium setup, loopless FVA, Pearson correlation), but only ~3 targets matched literature – likely due to not resetting objective to biomass before FVA under enforced ethanol scanning. Efficient for typical use; loopless FVA-in-loop may be slow. | **(Score: 42.75)** Similar to GPT but better readability/efficiency/error handling/documentation; same partial target match (~3) likely for same objective-reset issue. | **(Score: 39.75)** Implemented core logic + extra plotting, but missing loopless FBA, weaker context management/objective reset → no match to reported targets. | Not applicable (context window limitation) |
| FVSEOF with literature + GitHub code embedded | **(Score: 40)** Simplified the GitHub workflow; but substituted FBA for full FVA per reaction; blocked update of essential nutrients → only ~ 3 matches to literature targets. Note: GitHub vs literature target discrepancies exist but don't affect study validity. | **(Score: 43.25)** Similar to GitHub with improvements; cleaner structure; better biomass constraint handling; uses signed Pearson flux (more interpretable). Medium reset may be risky; used multiprocessing like GitHub but could leverage solver parallelism. | **(Score: 40.25)** Cleaner GitHub-based version; initial error due to regression failures from problematic numeric entries—fixable by filtering inf/NaN before stats. After correction, could reproduce GitHub results. | Not applicable |

**Table S10:** Sample prompt and response generated by Gemini upon comparing two GSMs.

**PROMPT:**

You are an expert in metabolic engineering, compare the iML1515 and yeast850 GSM models to cover the following:

1. Compare the model details?
2. what are the key metabolic reactions/pathways differences between the two models?

**RESPONSE:**
Of course. As an expert in metabolic engineering, here is a clear and concise comparison of the iML1515 and Yeast8.5.0 genome-scale metabolic models (GSMs).

The fundamental difference between these two models lies in the organisms they represent: iML1515 models the prokaryote ***Escherichia coli***, while Yeast8.5.0 models the eukaryote ***Saccharomyces cerevisiae***. This biological distinction drives nearly all the differences in their structure and metabolic capabilities.

### 1. Model Details Comparison

This table provides a high-level quantitative overview of the two models.

| Metric | iML1515 (*E. coli*) | Yeast8.5.0 (*S. cerevisiae*) | Key Takeaway |
| :--- | :--- | :--- | :--- |
| **Organism** | *Escherichia coli* K-12 MG1655 | *Saccharomyces cerevisiae* S288C | Prokaryote vs. Eukaryote |
| **Number of Genes** | 1,515 | 1,888 | Yeast8 is genetically more complex. |
| **Number of Reactions** | 2,712 | 4,058 | Yeast8 has significantly more reactions, largely due to compartmentalization. |
| **Number of Metabolites**| 1,877 | 2,742 | More metabolites in Yeast8, many being duplicates across different compartments. |
| **Compartments** | 3 (Cytoplasm, Periplasm, Extracellular) | 8 (Cytoplasm, Mitochondrion, Peroxisome, Nucleus, ER, Golgi, Vacuole, Extracellular) | **This is the most critical structural difference.** |

**In summary:** The Yeast8.5.0 model is substantially larger and more complex than iML1515. This increased complexity is a direct result of the eukaryotic cell structure of yeast, which requires the model to account for metabolites and reactions occurring in distinct organelles, as well as the transport mechanisms between them.

---

### 2. Key Metabolic Reactions/Pathways Differences

The differences in model details translate directly into significant differences in metabolic capabilities and network topology.

#### 1. Compartmentalization of Metabolism
This is the overarching difference that impacts all other pathways.
*   **iML1515 (*E. coli*):** Most metabolism, including glycolysis, the TCA cycle, and fatty acid synthesis, occurs in a single compartment (cytoplasm).
*   **Yeast8.5.0 (*S. cerevisiae*):** Metabolic pathways are distributed across organelles.
    *   **TCA Cycle:** Occurs in the **mitochondrion**. This requires the model to include numerous transporters for pyruvate, citrate, malate, etc., across the mitochondrial membrane.
    *   **Fatty Acid Oxidation (β-oxidation):** Occurs in both the **mitochondrion** and **peroxisome**.
    *   **Glyoxylate Shunt:** Key enzymes are located in the **cytoplasm** and **peroxisome**.

#### 2. Fermentation Pathways
This is a classic and critical difference for industrial biotechnology.
*   **iML1515 (*E. coli*):** Under anaerobic conditions, *E. coli* performs **mixed-acid fermentation**. The model includes pathways to produce lactate, formate, acetate, succinate, and ethanol, but ethanol is not the primary product.
*   **Yeast8.5.0 (*S. cerevisiae*):** Yeast is a premier **ethanol producer**. It possesses a highly efficient pathway using Pyruvate Decarboxylase (PDC) and Alcohol Dehydrogenase (ADH) to convert pyruvate almost exclusively to ethanol, even in the presence of oxygen (Crabtree effect). This pathway is a central feature of the Yeast8 model.

#### 3. Amino Acid Biosynthesis
While many pathways are conserved, there is a key difference in lysine synthesis.
*   **iML1515 (*E. coli*):** Uses the **diaminopimelate (DAP) pathway**.
*   **Yeast8.5.0 (*S. cerevisiae*):** Uses the **α-aminoadipate (AAA) pathway**. These pathways are completely distinct, involving different genes and intermediates.

#### 4. Lipid and Sterol Metabolism
This is a major point of divergence between prokaryotes and eukaryotes.
*   **iML1515 (*E. coli*):** Contains pathways for synthesizing fatty acids and phospholipids for its cell membrane. It **does not** have a pathway for sterol synthesis.

*   **Yeast8.5.0 (*S. cerevisiae*):** Contains an extensive and complex pathway for the synthesis of **ergosterol**, the primary sterol in the yeast membrane. This pathway, localized in the Endoplasmic Reticulum (ER), is a major metabolic sink and a common target for metabolic engineering to produce other high-value steroids and terpenoids.

#### 5. Protein Modification
High-quality GSMs include reactions for protein synthesis and modification.
*   **iML1515 (*E. coli*):** Includes reactions for protein synthesis and export to the periplasm but lacks complex post-translational modifications.
*   **Yeast8.5.0 (*S. cerevisiae*):** Includes reactions for **N-linked and O-linked glycosylation**, which occur in the ER and Golgi apparatus. This makes yeast a preferred host for producing complex eukaryotic proteins (e.g., pharmaceuticals) that require these modifications for proper function, a capability that is reflected in its model.

### Conclusion

The choice between using iML1515 or Yeast8.5.0 for an engineering project depends entirely on the biological objective.

*   **iML1515 (*E. coli*)** is ideal for modeling rapid growth and the production of many bulk chemicals, amino acids, and proteins that do not require complex modifications. Its relative simplicity makes simulations faster.
*   **Yeast8.5.0 (*S. cerevisiae*)** is essential for modeling processes that leverage unique eukaryotic features, such as efficient ethanol production, sterol-derived products, or the synthesis of complex glycoproteins. Its compartmentalization provides a more accurate representation of eukaryotic metabolism but adds significant computational complexity.

**Table S11:** GSM applications, limitations, and improvements/future directions provided by different LLMs.

| Category | Applications | Limitations | Improvements/Future directions |
|---|---|---|---|
| GPT | -Rational strain engineering using gene knockouts/overexpression<br>-OptKnock-guided strain design<br>-Pathway optimization with FBA<br>-High-value chemical production (e.g., 1.4-butanediol, isoprenoids) | -Incomplete/inaccurate network reconstructions<br>-Lack of regulatory and kinetic information<br>-Static/context-independent models<br>-Assumptions in objective functions reduce prediction accuracy | -Multi-omics integration<br>-Hybrid regulatory/kinetic model<br>-Machine learning integration<br>-Model standardization and Automated reconstruction tools (ModelSEED, CarveMe)<br>-Dynamic/spatial/community GSMs |
| Gemini | -Rational microbial cell factory design<br>-OptKnock and MOMA for gene knockout prediction<br>-Pathway optimization: bottleneck and cofactor balancing<br>-Production of BDO, artemisinic acid, lysine | -Incomplete genome annotations<br>-No enzyme kinetics or regulatory mechanisms<br>-Steady-state stoichiometric limitations<br>-Incorrect objective function assumptions | -Integration of transcriptomics, proteomics, metabolomics<br>-ME-models linking metabolism and gene expression<br>-ML/AI for gap-filling and regulation |

| | | | |
|---|---|---|---|
| | | | -Dynamic and whole-cell modelling |
| Claude | -Gene knockout strategies with OptKnock/MOMA<br>-FBA and FVA for pathway optimization<br>-Production of succinate, BDO, artemisinic acid, biofuels<br>-Cofactor balancing and precursor identification | -Steady-state assumption<br>-Missing regulatory information<br>-Biomass objective function<br>-Annotation gaps and transport uncertainty<br>-Multiple solution spaces and missing kinetic parameters | -Omics-constrained GSMs (GIMME, iMAT)<br>-Kinetic and regulatory integration<br>-Dynamic FBA<br>-ML-assisted flux prediction<br>-Whole-cell and community modelling<br>-Cloud and standardized platforms |
| DeepSeek | -Rational strain design and flux redirection<br>-OptFlux/OptStrain pathway optimization<br>-Dynamic FBA applications<br>-Production of succinate, lysine, taxadiene, artemisinin precursors | -Incomplete pathway annotations<br>-Neglect of transcriptional/translational regulation<br>-Steady-state assumption in FBA<br>-Missing enzyme kinetics, toxicity, compartmentalization effects | -Multi-omics + ML integration<br>-Dynamic and enzyme-constrained models (GECKO)<br>-Community-driven databases (BiGG. ModelSEED)<br>-Adaptive Laboratory Evolution (ALE)-guided refinement (validation and refine GSMs)<br>-Multi-scale and whole cell models<br>-AI-driven gap-filling and spatial modelling |

**Table S12:** Yield improvement strategies for limonene production suggested by different LLMs.

| LLM | Both hosts | iML1515 (E. coli) | Yeast850 | Distinctive features |
|---|---|---|---|---|
| GPT | Overexpress/codon-optimize limonene synthase; boost precursors IPP/DMAPP + GPP; reduce byproducts; cofactor (NADPH/ATP) balance; transport/efflux (reduce toxicity) | Add heterologous MVA or optimize native MEP; knockouts diverting flux from GPP/FPP | Upregulate HMG-CoA reductase (rate-limiting step); downregulate ERG9 (reduce FPP consumption); engineer GPP synthase | Emphasis on practical “metabolic engineering checklist” + transport to reduce toxicity |
| Gemini | Use FBA/OptKnock after adding LS reaction; boost NADPH via PPP | Overexpress dxs (DXP synthase), idi (IPP isomerase), and ispA (GPP synthase); knock out byproducts (ackA (acetate), ldhA (lactate), pflB (formate)); boost cofactor NADPH (MEP pathway) via zwf (glucose-6-phosphate dehydrogenase) | Overexpress tHMG1 (HMG-CoA reductase) + other MVA genes; engineer ERG20 for GPP; downregulate/KO ERG9 (prevent GPP/FPP diversion); redirect carbon to increase acetyl-CoA pool (MVA path) | Strongest on named gene targets + explicit computational workflow; adds compartment (e.g., mitochondria) targeting conceptually (distinct acetyl-CoA pools) |

| | | | | |
|---|---|---|---|---|
| Claude | MVA/MEP precursor boosting; efflux pumps; NADPH regeneration; knock out competing sinks; expression balancing | Introduce/optimize heterologous MVA; overexpress dxs, idi; knock out competing precursor drains | Upregulate HMG-CoA reductase; enhance acetyl CoA; optimize subcellular targeting | Adds adaptive laboratory evolution and lipid droplet sequestration for tolerance |
| DeepSeek | Cofactor engineering (NADPH) via PPP (pentose phosphate pathway) upregulation; secretion/efflux; in silico FBA/FVA; hybrid MVA/MEP; iterative validation | Overexpress dxs, idi, ispDF; feedback resistant enzymes; dynamic regulation (growth vs production); PPP upregulation | Overexpress tHMG1, ERG20; downregulate ERG9; boost cytosolic acetyl CoA (e.g., ACS2); block competing pathways (fatty acid synthesis) | Most explicit on dynamic regulation (decouple growth and production phases) + "host comparison" claim (yeast tolerance vs E. coli agility) |

**Table S13:** List of prompts and system prompts used for individual tasks.

| **Category: Domain knowledge** | | |
|---|---|---|
| **Task** | **Prompt** | **System prompt** |
| Model summarization and structure (JSON) | GSM model:<br>{json_data}<br><br>Given this iML1515.json GSM model,<br>1. summarize the details of the model?<br>2. provide the full structure in a visual tree? | You are an assistant who provides clear and concise responses to users. |
| Component identification (JSON) | GSM model:<br>{json_data}<br><br>Given this iML1515.json GSM model,<br>1. list all reactions involving acetyl-CoA as a substrate in a table,<br>2. and summarize the involved pathways? | You are an assistant who provides clear and concise responses to users. |
| Component identification (JSON) using code | GSM model:<br>{json_data}<br><br>Given this iML1515.json GSM model,<br>1. list all reactions involving acetyl-CoA as a substrate in a table,<br>2. and summarize the involved pathways? | You are an assistant who provides clear and concise responses to users.\<br>Provide full python code first, then only followed by step-by-step explanations. \<br>When providing python code, starting with ```python and end with ```.\<br>Please think properly to ensure that the code can be run properly with high efficiency. |
| Subsystem identification (JSON) | Read this iML1515.json GSM model:<br>{json_data}<br><br>Summarize the reactions or pathways involving NAD+/NADH or NADP+/NADPH? | You are an assistant who provides clear and concise responses to users. |
| Biomass composition (JSON) | Read this iML1515.json GSM model:<br>{json_data} | You are an assistant who provides clear and concise responses to users. |

| | | |
|---|---|---|
| | 1. Explain the composition and role of the biomass objective function in the model?<br>2. What are the key differences between WT and core biomass? | |
| Pathway prediction, host selection | Propose a biosynthetic pathway to produce limonene in a microbial host using the mevalonate (MVA) pathway.<br>1. List the necessary genes and enzymes involved from acetyl-CoA to limonene, including intermediate metabolites and any required heterologous enzymes.<br>2. Suggest possible host organisms,<br>3. Highlight key regulatory points or potential bottlenecks in the pathway.<br>4. Recommend genetic modifications to enhance limonene yield. | You are an assistant who provides clear and concise responses to users. |
| GSM models comparison (*truncated JSON) | Read and compare these two iML1515 and yeast850 GSM models:<br>iML1515 model:<br>{iML1515_info}<br><br>yeast850 model:<br>{yeast850_info}<br><br>1. Based on the provided data, what are the key metabolic reactions/pathways differences between the two models? | You are an assistant who provides clear and concise responses to users. |
| GSM models comparison | You are an expert in metabolic engineering, compare the iML1515 and yeast850 GSM models to cover the following:<br><br>1. Compare the model details?<br>2. what are the key metabolic reactions/pathways differences between the two models? | You are an assistant who provides clear and concise responses to users. |
| GSM applications, limitations, improvements | You are an expert in systems biology and metabolic engineering, provide a detailed analysis of the role of GSM covering the following points:<br><br>1. Applications: How GSMs are applied in strain design, pathway optimization, and bioproduction of high-value chemicals. Provide successful examples.<br>2. Limitations: Highlight the key challenges with GSM, and how these limitations affect prediction accuracy.<br>3. Improvements: Opportunities for improving GSM predictive power, and future directions. | You are an assistant who provides clear and concise responses to users. |
| Growth-coupled concepts | You are an expert in systems biology and metabolic engineering, answer the following points:<br><br>1. Definition: Explain what it means for a metabolic pathway or product to be growth-coupled. Contrast weak vs strong growth coupling.<br>2. Analysis: How GSM model can be used to determine if a pathway is growth-coupled? suggest tools or methods and compare. | You are an assistant who provides clear and concise responses to users. |

| | |
|---|---|
| | 3. Engineering Relevance: why is growth-coupling important in metabolic engineering and strain design? support with successful examples.<br>4. Limitations and challenges: What are the limitations of using in silico growth-coupling analysis? How can experiment observations diverge from model predictions.<br>5. Future directions: Opportunities for improving growth-coupled strategies computationally and experimentally? |

| **Category: Flux Prediction** | | |
|---|---|---|
| **Task** | **Prompt** | **System prompt** |
| Run FBA analysis | provide python code to run FBA analysis for iML1515.json GSM model with explanations, and print the result and suggest top 10 reactions with the highest flux? | You are an assistant who provides clear and concise responses to users.\ Provide full python code first, then only followed by step-by-step explanations. \ When providing python code, starting with ```python and end with ```.\ Please think properly to ensure that the code can be run properly with high efficiency. |
| Run FVA analysis | provide python code to run FVA analysis for iML1515.json GSM model with explanations, print the result and identify the top 10 redundant reactions with the largest flux range? | You are an assistant who provides clear and concise responses to users.\ Provide full python code first, then only followed by step-by-step explanations. \ When providing python code, starting with ```python and end with ```.\ Please think properly to ensure that the code can be run properly with high efficiency. |

| **Category: Metabolic pathway construction** | | |
|---|---|---|
| **Task** | **Prompt** | **System prompt** |
| Gap identification for pathway (JSON) | Read this iML1515.json GSM model:<br>{json_data}<br><br>Suggest candidate reactions and enzymes that could fill metabolic gaps to allow for compound limonene synthesis? | You are an assistant who provides clear and concise responses to users. |
| GSM model comparison for pathway | You are an expert in metabolic engineering, compare the iML1515 and yeast850 GSM models for limonene biosynthesis by covering the following:<br><br>1. What are the metabolic gaps for using these models for limonene biosynthesis?<br>2. Which model is better for limonene biosynthesis? | You are an assistant who provides clear and concise responses to users. |

| | | |
|---|---|---|
| | 3. How to improve the limonene yield in these models. | |
| Pathway integration into GSM, run FBA | Propose a biosynthetic pathway to produce limonene in a microbial host using the mevalonate (MVA) pathway. \ 1. List the necessary genes and enzymes involved from acetyl-CoA to limonene, including intermediate metabolites and any required heterologous enzymes. \ 2. provide python script to add this pathway into iML1515.json GSM model, and run FBA analysis to optimize limonene. | You are an assistant who provides clear and concise responses to users.\ Provide full python code first, then only followed by step-by-step explanations. \ When providing python code, starting with ```python and end with ```.\ Please think properly to ensure that the code can be run properly with high efficiency. |
| **Category: Metabolic flux optimization** | | |
| **Task** | **Prompt** | **System prompt** |
| FVSEOF for succinate in E. coli iML1515 | Provide Python code to run FVSEOF for the iML1515.json GSM model to improve succinate production. At each enforced succinate production level, use Flux Variability Analysis (FVA) to obtain minimum and maximum fluxes for all reactions. Then, for each reaction, calculate the correlation between the enforced succinate level and the minimum flux value (and optionally, maximum flux value). Identify and report the top 10 reactions whose minimum flux is most positively correlated with succinate (upregulation targets), and the top 10 most negatively correlated (downregulation targets). Please include clear comments in the code and explanations. | You are an assistant who provides clear and concise responses to users.\ Provide full python code first, then only followed by step-by-step explanations. \ When providing python code, starting with ```python and end with ```.\ Please think properly to ensure that the code can be run properly with high efficiency. |
| Single-knockout analysis to improve product yield (e.g., limonene) while maintaining 10% max biomass | We have a iML1515_limonene.json GSM file with limonene pathway through the mevalonate (MVA) pathway, \ 1. provide python script to run single-knockout analysis using cobrapy to improve limonene yield, while maintaining 10% of max biomass. | You are an assistant who provides clear and concise responses to users.\ Provide full python code first, then only followed by step-by-step explanations. \ When providing python code, starting with ```python and end with ```.\ Please think properly to ensure that the code can be run properly with high efficiency. |
| Succinate knockout workflow in E. coli iJR904 | You are an expert in metabolic engineering, \ 1. suggest genes to knock-out to improve succinate production in iJR904 GSM model at anaerobic condition.\ 2. provide python script to include the following:\ a. run FBA to identify succinate flux when optimizing biomass at anaerobic condition.\ b. run FBA to optimize succinate at anaerobic condition while maintaining 10% of wild-type biomass flux.\ | You are an assistant who provides clear and concise responses to users.\ Provide full python code first, then only followed by step-by-step explanations. \ When providing python code, starting with ```python and end with ```.\ Please think properly to ensure that the code can be run properly with high efficiency. |

| | | |
|---|---|---|
| | c. knockout the individual suggested genes using cobrapy package to determine the impacts on yield and biomass. | |
| Growth-coupled analysis (e.g., limonene pathway in iML1515) | We have a iML1515_limonene.json GSM file with limonene pathway through the mevalonate (MVA) pathway, \ <br> 1. provide python script using cobrapy to determine if the limonene production is growth-coupled. | You are an assistant who provides clear and concise responses to users.\ <br> Provide full python code first, then only followed by step-by-step explanations. \ <br> When providing python code, starting with ```python and end with ```.\ <br> Please think properly to ensure that the code can be run properly with high efficiency. |

**Category: Case study validation**

| Task | Prompt | System prompt |
|---|---|---|
| Read and summarize literature | Read and summarize the given paper: <br><br> Paper: <br> {Literature1} | You are an assistant who provides clear and concise responses to users. |
| Compute YT & YA for L-lysine in *E. coli* GSM; D-glucose, aerobic | Paper: <br> {Literature1} <br><br> Based on this paper, provide python code using Cobrapy to compute the theoretical yield and maximum achievable yield of L-lysine in *E. coli* iML1515.xml GSM model using D-glucose as carbon source under aerobic condition. | You are an assistant who provides clear and concise responses to users.\ <br> Provide full python code first, then only followed by step-by-step explanations. \ <br> When providing python code, starting with ```python and end with ```.\ <br> Please think properly to ensure that the code can be run properly with high efficiency. |
| Compute YT & YA for ethanol in *E. coli* under aerobic/anaerobic/microaerobic conditions | Paper: <br> {Literature1} <br><br> Based on this paper, provide python code using Cobrapy to cover the following: <br> 1. to compute the theoretical yield and maximum achievable yield of ethanol in *E. coli* iML1515.xml GSM model using D-glucose as carbon source under aerobic condition, <br> 2. under anaerobic condition, <br> 3. under microaerobic condition using the setting used in the paper. <br> *Note: use the default setting of the medium except for D-glucose. | You are an assistant who provides clear and concise responses to users.\ <br> Provide full python code first, then only followed by step-by-step explanations. \ <br> When providing python code, starting with ```python and end with ```.\ <br> Please think properly to ensure that the code can be run properly with high efficiency. |
| Compute YT & YA for ethanol in Yeast under aerobic/anaerobic/microaerobic conditions | Paper: <br> {Literature1} <br><br> Based on this paper, provide python code using Cobrapy to cover the following: <br> 1. to compute the theoretical yield and maximum achievable yield of ethanol in S. cerevisiae | You are an assistant who provides clear and concise responses to users.\ <br> Provide full python code first, then only followed by step-by-step explanations. \ |

| | | |
|---|---|---|
| | yeast850.xml GSM model using D-glucose as carbon source under aerobic condition,<br>2. under anaerobic condition,<br>3. under microaerobic condition using the setting used in the paper.<br>*Note: use the default setting of the medium except for D-glucose. | When providing python code, starting with ```python and end with ```.\ Please think properly to ensure that the code can be run properly with high efficiency. |
| Pathway integration into C. glutamicum GSM: add R-mevalonate + FBA for YT | iCW773 GSM model:<br>{iCW773_json}<br><br>Pathway:<br>{pathway}<br><br>Read the iCW773.json model, provide python code using Cobrapy to cover the following:<br>1. to add the R-mevalonate production pathway into the iCW773.json GSM model,<br>2. to run FBA analysis to determine the YT molar yield (mol/mol) using D-glucose as carbon source under aerobic condition,<br>*To calculate YT theoretical yield, the lower bound of ATPM reaction was set to zero.<br>*D-glucose update flux = -10; O2 = -1000;<br>*Note: use the default setting of the medium provided by the model except for D-glucose.<br>*Note: Check the internal metabolites to avoid duplicates when adding pathway. | You are an assistant who provides clear and concise responses to users.\ Provide full python code first, then only followed by step-by-step explanations. \ When providing python code, starting with ```python and end with ```.\ Please think properly to ensure that the code can be run properly with high efficiency. |
| Implement iBridge algorithm based on literature | Paper:<br>{Literature1}<br><br>Based on this paper, provide python code using Cobrapy to cover the following:<br>1. to run iBridge to identify up- and down-regulation targets to improve the production flux of ethanol in *E. coli* iML1515.xml GSM model using D-glucose as carbon source under aerobic condition.<br><br>*Note: use the default setting of the medium provided by the model except for D-glucose. | You are an assistant who provides clear and concise responses to users.\ Provide full python code first, then only followed by step-by-step explanations. \ When providing python code, starting with ```python and end with ```.\ Please think properly to ensure that the code can be run properly with high efficiency. |
| Implement iBridge with literature + GitHub code embedded | Paper:<br>{Literature1}<br><br>GitHub code:<br>{code}<br><br>Based on this paper and the provided code, provide python code using Cobrapy to cover the following:<br>1. to run iBridge to identify up- and down-regulation targets to improve the production flux of ethanol in *E. coli* iML1515.xml GSM model using D-glucose as carbon source under aerobic condition.<br>2. by considering the excluded metabolites provided in the excluded_metabolites.txt. | You are an assistant who provides clear and concise responses to users.\ Provide full python code first, then only followed by step-by-step explanations. \ When providing python code, starting with ```python and end with ```.\ Please think properly to ensure that the code can be run properly with high efficiency. |

| | | |
|---|---|---|
| | *Note: use the default setting of the medium provided by the model except for D-glucose. | |
| FVSEOF from literature only | Paper:<br>{Literature1}<br><br>Based on the descriptions provided in this paper, provide python code using Cobrapy to cover the following:<br>1. to run FVSEOF to identify up-regulation targets to improve the production flux of ethanol in *E. coli* iML1515.xml GSM model using D-glucose as carbon source under aerobic condition.<br><br>*Note: use the default setting of the medium provided by the model except for D-glucose. | You are an assistant who provides clear and concise responses to users.\<br>Provide full python code first, then only followed by step-by-step explanations. \<br>When providing python code, starting with ```python and end with ```.\<br>Please think properly to ensure that the code can be run properly with high efficiency. |
| FVSEOF with literature + GitHub code embedded | Paper:<br>{Literature1}<br><br>GitHub code:<br>{code}<br><br>Based on the descriptions provided in this paper and the provided code, provide python code using Cobrapy to cover the following:<br>1. to run FVSEOF to identify up-regulation targets to improve the production flux of ethanol in *E. coli* iML1515.xml GSM model using D-glucose as carbon source under aerobic condition.<br><br>*Note: use the default setting of the medium provided by the model except for D-glucose. | You are an assistant who provides clear and concise responses to users.\<br>Provide full python code first, then only followed by step-by-step explanations. \<br>When providing python code, starting with ```python and end with ```.\<br>Please think properly to ensure that the code can be run properly with high efficiency. |

**Table S14:** Analysis strategies used in evaluating LLM error-detection capability

| **(i) Detect stoichiometric sign error:** | | |
|---|---|---|
| **Task** | **Prompt** | **System Prompt** |
| (text-based interpretation using general prompting) | Read this iML1515_case1.json GSM model:<br>{json_data}<br><br>identify if there is any stoichiometry imbalance issue in the model? | You are an assistant who provides clear and concise responses to users. |

| | | |
|---|---|---|
| (text-based interpretation using general prompting) | Read this iML1515_case2.json GSM model:<br>{json_data}<br>identify if there is any issue in the model? | You are an assistant who provides clear and concise responses to users. |
| (text-based interpretation using general prompting) | Read this iML1515_case2.json GSM model:<br>{json_data}<br><br>identify if there is any missing reaction in the model? | You are an assistant who provides clear and concise responses to users. |
| (code-based analysis using general prompting) | Read this iML1515_case2.json GSM model:<br>{json_data}<br><br>identify if there is any missing reaction in the model? | You are an assistant who provides clear and concise responses to users. \<br>if you need to provide code, provide full python code first, then only followed by step-by-step explanations. \<br>When providing python code, starting with ```python and end with ```.\<br>Please think properly to ensure that the code can be run properly with high efficiency |
| (text-based interpretation using specific prompting with guided steps) | Read this iML1515_case2.json GSM model:<br>{json_data}<br><br>Perform a pathway-completeness analysis of central carbon metabolism, especially glycolysis.<br><br>Your task:<br>1. List the canonical glycolysis reactions expected in E. coli.<br>2. Check whether each expected reaction is present in the model.<br>3. Identify any missing internal reaction.<br>4. Provide the expected reaction ID, name, equation, and affected metabolites.<br>5. Explain the metabolic consequence of the missing reaction.<br>6. Ignore exchange, demand, sink, and biomass pseudo-reactions when deciding whether a gap exists.<br><br>Return a concise but evidence-based answer. | You are an assistant who provides clear and concise responses to users. |
| (code-based cross-checking against reference model) | You are given two genome-scale metabolic models in COBRA JSON format.<br><br>Reference model:<br>{original_json_data}<br><br>Test model:<br>{json_data}<br><br>Your task is to use code-based analysis to compare the reference model against the test | You are an assistant who provides clear and concise responses to users. \<br>if you need to provide code, provide full python code first, then only followed by step-by-step explanations. \<br>When providing python code, starting with ```python and end with ```.\ |

| | | |
|---|---|---|
| | model and identify any missing **internal reactions** or lost metabolic connectivity. | Please think properly to ensure that the code can be run properly with high efficiency |
| (code-based cross-checking against reference model) | You are given two genome-scale metabolic models in COBRA JSON format.<br><br>Reference model:<br>{original_json_data}<br><br>Test model:<br>{json_data}<br><br>identify if there are any differences between the reference model and the test model? | You are an assistant who provides clear and concise responses to users. \ <br>if you need to provide code, provide full python code first, then only followed by step-by-step explanations. \ <br>When providing python code, starting with ```python and end with ```.\ <br>Please think properly to ensure that the code can be run properly with high efficiency |
| (text-based cross-checking against reference model) | You are given two genome-scale metabolic models in COBRA JSON format.<br><br>Reference model:<br>{original_json_data}<br><br>Test model:<br>{json_data}<br><br>identify if there are any differences between the reference model and the test model? | You are an assistant who provides clear and concise responses to users. |
